# Revisiting flippase specificity: Drs2-Cdc50 transports multiple anionic lipid substrates

**DOI:** 10.1101/2025.08.22.671827

**Authors:** Amalie B. Purup, Pauline Funke, Cédric Montigny, Aurélie Di Cicco, Thibaud Dieudonné, Merethe M. Frøsig, Yugo Iwasaki, Daniel Lévy, Thomas Günther Pomorski, Rosa L. López-Marqués, Joseph A. Lyons, Guillaume Lenoir

## Abstract

P4-ATPase lipid flippases maintain transbilayer lipid asymmetry in eukaryotic membranes, which is essential for many cellular processes. In yeast, the Drs2-Cdc50 flippase complex was previously shown to specifically transport phosphatidylserine (PS) from the exoplasmic to the cytosolic leaflet of the *trans*-Golgi network (TGN), thereby controlling vesicular trafficking in the secretory and endocytic pathways. Using an improved proteoliposome-based lipid flippase assay, we now show that the Drs2-Cdc50 complex transports multiple anionic glycerophospholipids, including PS, phosphatidylinositol, phosphatidylglycerol, and phosphatidic acid. *In vivo* cell-based lipid uptake assays further support the transport of these lipids. To understand the basis of this substrate promiscuity, we analyzed cryo-EM structures of the complex with occluded lipids. These structures revealed that the water network surrounding the lipid headgroup plays a critical role in enabling Drs2-Cdc50 to recognize different lipids. These data unveil an unexpected broad specificity of the Drs2-Cdc50 complex for anionic lipids, which may significantly impact their transbilayer distribution in the yeast TGN.

## Introduction

A striking feature of eukaryotic cell membranes is the asymmetric distribution of lipids across the bilayer (*1*). In membranes of the late secretory/endocytic pathways, including the plasma membrane, the endosomes and the *trans*-Golgi network (TGN), sphingolipids are confined to the exoplasmic leaflet, while phosphatidylserine (PS), phosphatidylethanolamine (PE), phosphatidylinositol (PI) and phosphoinositides (PIPs) are largely restricted to the cytosolic leaflet (*2–5*). Such transbilayer lipid asymmetry fulfills a host of cellular functions. For example, the uneven distribution of PS, PI and PIPs results in a high negative surface charge in the late endomembrane system, which is used to regulate the recruitment of peripheral membrane proteins through their polybasic motifs (*6*, *7*).

To maintain lipid asymmetry, cells have evolved lipid transporters, termed flippases, which actively transport lipids toward the cytosolic leaflet of cell membranes. Most eukaryotic flippases belong to the P4 subfamily of P-type ATPases (P4-ATPases), a family of polytopic membrane proteins that catalyze substrate transport at the expense of ATP hydrolysis (*8*). Importantly, most mature flippases consist of a complex formed by a P4-ATPase and a CDC50 subunit, the latter being essential for the proper subcellular localization and catalytic function of P4-ATPases (*9–11*). From a structural standpoint, P4-ATPases are composed of ten transmembrane spans that harbor the lipid binding cavity, and three cytosolic domains that bind and hydrolyze ATP (*12–15*). The N-domain binds ATP, the P-domain is phosphorylated from ATP on a conserved aspartate residue, while the A-domain resets the enzyme to its initial state by catalyzing dephosphorylation of the P-domain. Phosphorylation/dephosphorylation of P4-ATPases is tightly coupled to conformational changes in the membrane domain, thereby allowing alternating access of the transported lipid to each side of the membrane (*16*). Collectively, the various states adopted by P4-ATPases are described in a reaction cycle known as the Post-Albers cycle (fig. S1). The physiological relevance of P4-ATPases is underscored by the fact that mutations in these proteins are associated with pathological conditions. For example, mutations in the human ATP8A2 and ATP8B1 genes result in neurological disorders and hepatic disorders, respectively (*17*, *18*). Meanwhile, variants of ATP8B4 have been identified as risk factors for systemic sclerosis and Alzheimer’s disease (*19–21*).

The lipid substrate specificity of several P4-ATPases has been addressed primarily by monitoring the uptake of fluorescent lipids in cell-based assays. P4-ATPases transport a variety of lipid substrates, including the major glycerophospholipids (PS, PE, and phosphatidylcholine (PC)), signaling lysoglycerophospholipids (e.g. lyso-PC) and ceramide-derived lipids such as glucosyl- and galactosylceramide. However, most P4-ATPases recognize specific lipids (mainly PC, PE or PS) and only a few P4-ATPases have been demonstrated to recognize a wide variety of transport substrates (*22*). Mammalian ATP8A2, was shown to be specific for PS after reconstitution in proteoliposomes (*23*). Likewise, ATP11A, ATP11B and ATP11C flip PS, and to a lesser extent PE (*24*). On the contrary, plant ALA10 was shown to recognize PC, lyso-PC, PS, PE, PG, PA, and even sphingomyelin, a mammalian lipid (*25*). Interestingly, recent studies have pointed out novel putative substrates for P4-ATPases, such as PI (*26–28*). For instance, ATP8B1, ATP8B2 and ATP10A, which were initially characterized as PC transporters, were recently shown to transport PI using cell-based transport assays (*26*). However, conclusive evidence of PI recognition and the broad substrate specificity of some P4-ATPases requires reconstitution of the purified transporters in proteoliposomes to avoid interference from other cellular components. Such an isolated system would be instrumental to understand the precise molecular mechanism by which P4-ATPases recognize various lipid species.

The yeast Drs2-Cdc50 lipid flippase complex is a founding member of the P4-ATPase family (*29*). The three-dimensional structure of Drs2-Cdc50 was among the first to be determined, revealing the tight regulation of the complex (*12*, *14*, *30*). We and others previously showed that the complex is autoinhibited by its N- and C-terminal tails (*12*, *14*, *31*) while on the other hand, its activity depends strongly on the presence of phosphatidylinositol-4-phosphate (PI4P), a phosphoinositide that controls vesicle trafficking at TGN membranes (*31–33*). The Drs2-Cdc50 complex is localized in membranes of the TGN, a transition point for the establishment of lipid asymmetry (*1*). Drs2-Cdc50 has been shown to transport NBD-PS using purified Golgi membranes, but not NBD-PC or NBD-PE, thereby maintaining PS asymmetry in the TGN (*34*). Moreover, proteoliposomes reconstituted with Drs2-Cdc50 display NBD-PS flippase activity (*35*, *36*). Drs2-Cdc50 also plays an essential role in membrane trafficking events, where its flippase activity is required for the formation of clathrin-coated vesicles from the TGN and for bidirectional transport between the TGN and early endosomes, suggesting that PS transport supports vesicle formation at the TGN (*37*, *38*). However, the involvement of Drs2 in vesicle trafficking is strikingly independent of the presence of PS, suggesting that other lipid substrates may fulfill this role (*34*).

Using improved biochemical, cell-based, and structural approaches, we now report broad anionic lipid specificity for the Drs2-Cdc50 complex. Specifically, our data indicates that the Drs2-Cdc50 ATPase activity is stimulated by a wide range of anionic phospholipids, including unlabeled PS, PI, PG, and PA. Upon reconstitution of the purified complex in proteoliposomes, we found that NBD-labeled derivatives of these negatively charged lipids are transported across the liposomal membrane in a PI4P-dependent manner, provided autoinhibitory Drs2 extensions are removed. Transport of these lipids was also confirmed by Drs2-Cdc50 overexpression in yeast mutants lacking endogenous P4-ATPases. To further explore the molecular basis of these findings, we performed cryo-electron microscopy (cryo-EM) analysis of the Drs2-Cdc50 complex. Our approach focused on the binding and occlusion of Drs2-Cdc50 lipid substrates, providing mechanistic insights into substrate recognition and selectivity. Together, our results identify a mechanism by which anionic lipids, including PI, remain asymmetric in TGN membranes and call for a re-evaluation of the substrate specificity of other P4-ATPases.

## Results

### Purified Drs2-Cdc50 complex displays ATPase activity towards anionic phospholipids

*In vitro* substrate-stimulated ATPase activity of P-type ATPases is often used to screen potential transport substrates and activators (*39–41*). To assess substrate specificity of Drs2-Cdc50, we examined its *in vitro* ATPase activity in response to various phospholipids. We expressed and purified three Drs2 constructs: Full-length wild-type (WT) Drs2 (FL Drs2), which is autoinhibited by its N- and C-terminal extensions, a WT Drs2 construct with cleavable N- and C-termini (ΔNC Drs2, comprising residues 104-1247), and an ATPase-deficient Drs2 mutant (ΔNC Drs2^E342Q^), also with cleavable N- and C-termini, locked in the E2P state, i.e. unable to complete a full catalytic cycle (fig. S1, fig. S2A). Truncation of both N- and C-terminal extensions, together with the presence of PI4P, is required to activate Drs2 *in vitro* (*31*). Affinity purification via a biotin acceptor domain tag followed by size-exclusion chromatography (SEC) yielded pure FL Drs2, ΔNC Drs2 and ΔNC Drs2^E342Q^ in complex with Cdc50 (fig. S2B). To confirm the functionality of the purified protein complexes, we measured their ATPase activity in the presence of PS and PI4P. Detergent-solubilized ΔNC Drs2 showed robust activity, which was inhibited by beryllium fluoride (BeFx), a general inhibitor of P-type ATPases that acts as a phosphate mimic, while ΔNC Drs2^E342Q^ exhibited no activity, as expected (fig. S2C). FL Drs2 remained largely inactive, even in the presence of PS and PI4P, but became active following enzymatic cleavage of its N- and C-terminal extensions by trypsin, consistent with previous studies (*31*) (fig. S2D).

We next screened for lipid specificity by measuring ATPase activity of detergent-solubilized, SEC-purified ΔNC Drs2 in the presence of PI4P and various natural phospholipids. Surprisingly, while PS is expected to be the primary substrate for Drs2, we observed a broad anionic lipid-stimulated ATPase activity including PS, PI, phosphatidylglycerol (PG), and, to a lesser extent, phosphatidic acid (PA), whereas PC and PE had no significant effect (Fig. 1A).

**Fig. 1:**
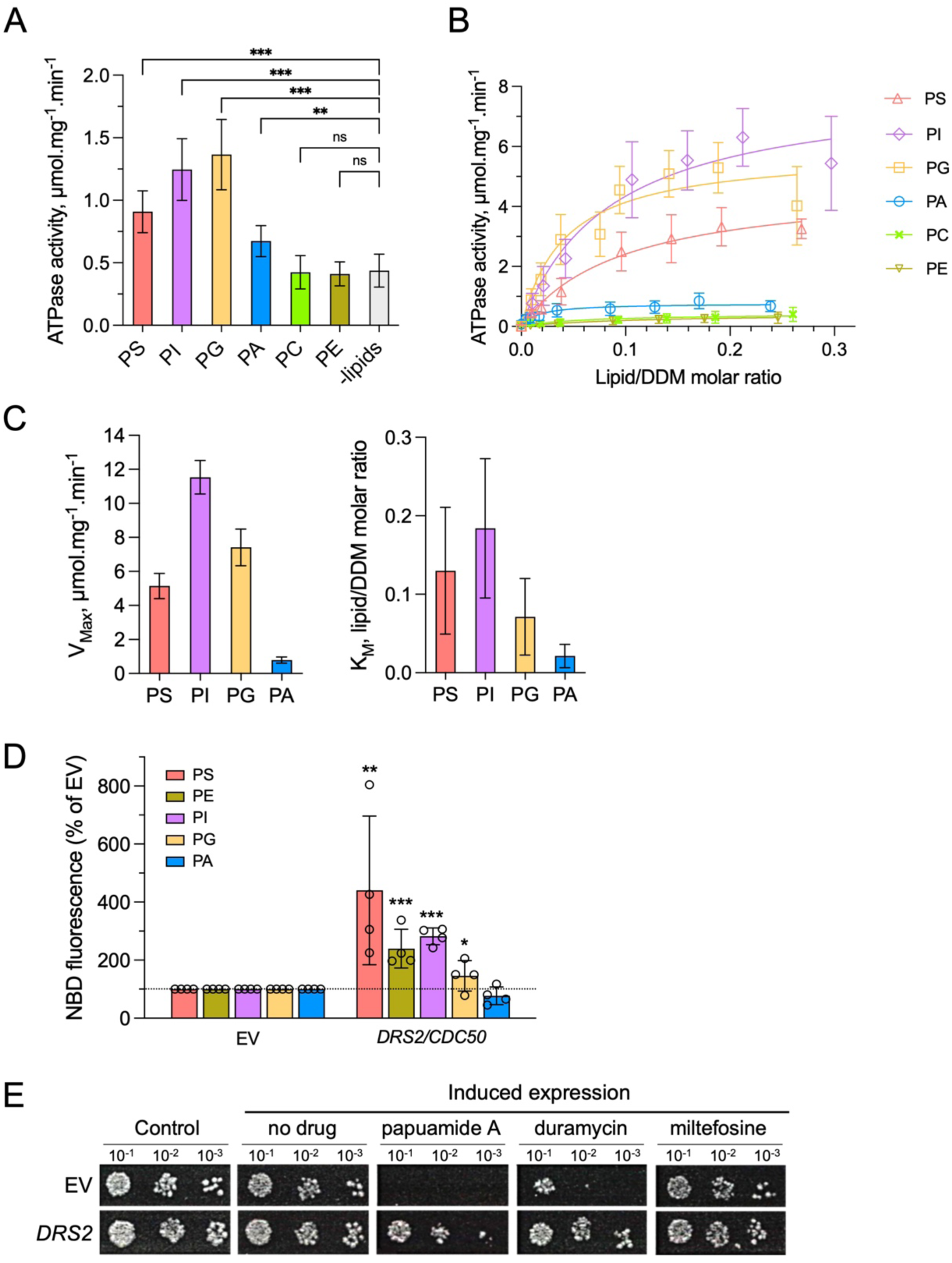
Lipid-stimulated ATPase activity and Drs2-mediated lipid transport *in vivo*. **(A)**, Specific ATPase activity of detergent-solubilized ΔNC Drs2-Cdc50 (10 µg/mL) in the presence of DDM (1 mg/mL), PI4P (25 µg/mL) and the indicated phospholipids (6.25 µg/mL, corresponding to a lipid/DDM molar ratio of ∼0.01), measured via a phosphomolybdate assay. Bars show the mean ± SD (n = 3-7); significance by one-way ANOVA (***p≤0.001, **p≤0.01, ns p>0.01). **(B)**, Specific ATPase activity of ΔNC Drs2-Cdc50 across lipid/DDM molar ratios. Data (mean ± SD, n= 3-4) were baseline-corrected and fitted to the Michaelis-Menten equation (y=v_max_ •x/(K_M_+x)). **(C)**, Michaelis-Menten constant (K_M_) and maximal velocity (V_max_) obtained from the fit in panel B. Bars represent the mean ± SD of 3 independent fits. PE and PC were excluded due to poor fit. **(D)**, Yeast *Δdrs2dnf1dnf2* mutant cells expressing Drs2 and Cdc50 or transformed with an empty vector (EV) were incubated with NBD-lipids and fluorescence intensity was measured by flow cytometry. Accumulation of NBD lipids is expressed as percentage of fluorescence intensity relative to the EV control (dashed line indicates 100%). Results are averages ± sem of 4 independent experiments for all samples except NBD-PG (n=3). One hundred percent corresponds to 4.68 ± 0.29 arbitrary units (NBD-PI), 5.96 ± 0.42 arbitrary units (NBD-PS), 7.11 ± 0.11 arbitrary units (NBD-PE), 5.45 ± 0.46 arbitrary units (NBD-PA), and 8.04 ± 0.65 arbitrary units (NBD-PG). **(E)**, A *Δdrs2dnf1dnf2dnf3* yeast mutant was transformed with an empty vector or a plasmid expressing *DRS2* under the control of a galactose-inducible promoter. Cells were dropped to the indicated OD_600_ on control plates with glucose or on galactose plates (induced expression) with no further additions (no drug) or supplemented with 0.1 µg/mL papuamide A, 1.5 µM duramycin, or 2.5 µg/mL miltefosine, as indicated. The figure shows a representative experiment from 3 independent repetitions. *p<0.05, **p<0.01, ***p<0.001 in a paired one-tailed t-test assuming equal variances.

To quantify lipid preference, we analyzed ATPase activity across increasing lipid/DDM molar ratios (0–0.3). Activity increased with lipid concentration, particularly for PI, PG, and PS, although saturation was not reached (Fig. 1B). Michaelis-Menten analysis revealed similar K_m_ values for PS, PI, PG, and PA, indicating comparable affinities, while V_max_ values differed, with PA yielding the lowest turnover (Fig. 1 C). Notably, neither PC nor PE stimulated activity, in contrast with previous claims that PE is a substrate (*42*, *43*). Collectively, these findings suggest broader specificity for anionic phospholipids, including PI, PG, and PA - substrates not previously associated with Drs2-Cdc50.

### Drs2 transports anionic phospholipids in living cells

To further substantiate the broad specificity of Drs2-Cdc50 for anionic lipids, the flippase complex was overexpressed in *Δdrs2dnf1dnf2* mutant yeast cells and the lipid transport activity was tested using lipid analogs labeled with a fluorescent NBD moiety on the *sn-2* acyl chain (*44*). Due to the lack of three endogenous flippases, namely Drs2, Dnf1 and Dnf2, the *Δdrs2dnf1dn2* mutant has a low level of endogenous lipid transport across the plasma membrane and has been extensively used for assigning the specificity of lipid flippases (*25*, *45–47*). As expected, *Δdrs2dnf1dnf2* cells overexpressing the Drs2-Cdc50 complex showed clear NBD-PS transport, about 4.5-fold higher than that of the empty vector control (Fig. 1D). The transport of NBD-PI and NBD-PE was about three times that of the empty vector, while the fluorescent signal for NBD-PG increased by 1.7-fold. NBD-PA uptake was not observed, probably due to technical limitations of the assay derived from the small PA head group (*48*). These data confirm that Drs2 is a broad-specificity transporter recognizing several anionic phospholipids, including PS, PI and PG.

Transport of NBD-PE does not correlate with the lack of PE-stimulated ATPase activity for purified Drs2. To verify whether natural PE is a substrate of Drs2, we employed a quadruple yeast mutant lacking all endogenous flippases except the essential Neo1 (*Δdrs2dnf1dnf2dnf3*). A wild-type yeast strain has an asymmetric distribution of phospholipids across the plasma membrane with PS and PE concentrated in the inner leaflet. This asymmetry can be probed with cytotoxic peptides that bind to PS (papuamide A) or PE (duramycin) when they are exposed on the cell surface. As expected, the quadruple *Δdrs2dnf1dnf2dnf3* mutant cannot survive in the presence of relatively low amounts of papuamide A and duramycin (Fig. 1E, empty vector control). Expression of *DRS2* from a plasmid resulted in yeast cells surviving on plates containing the cytotoxic peptides (Fig. 1E), indicating recovery of the PS and PE asymmetry. In line with the inability of Drs2 to recognize the phosphocholine headgroup, *Δdrs2dnf1dnf2dnf3* cells expressing *DRS2* can survive in the presence of miltefosine, a lyso-PC analog that is toxic when taken up by plasma membrane flippases. These results suggest that Drs2 might transport natural PE, despite the inability of this lipid to stimulate ATPase activity *in vitro*.

### Drs2-Cdc50 actively transports anionic phospholipids in reconstituted vesicles

To test whether Drs2 can translocate lipids beyond PS, as indicated by the ATPase activity and in-cell lipid uptake measurements, we used a dithionite-based fluorescence quenching assay (Fig. 2A). While effective for studying lipid scramblases (e.g. (*49*, *50*)), this assay is more challenging for flippases due to the unidirectional transport they catalyze, which may impose asymmetrical stress on the bilayer (*51*, *52*). Nonetheless, prior studies have confirmed Drs2-mediated PS transport, though its broader substrate specificity remained uncharacterized (*35*, *36*).

**Fig. 2:**
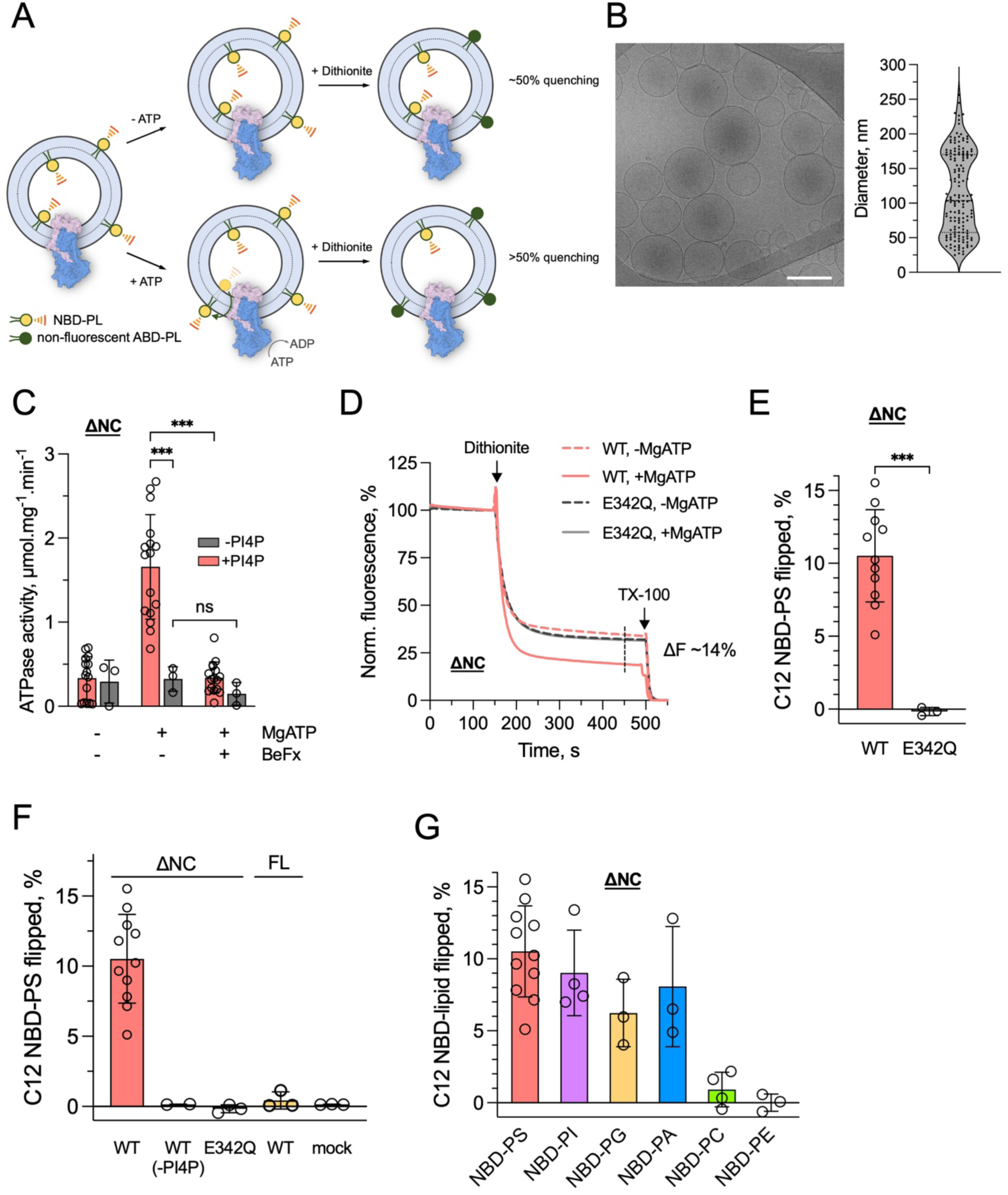
Drs2-mediated lipid transport in proteoliposomes. **(A)**, Schematic of the fluorescence-based lipid transport assay. The reconstituted ΔNC Drs2-Cdc50 complex with the catalytic domains oriented towards the assay medium actively flips NBD-lipids from the inside to the outside of the vesicle only in the presence of MgATP, whereas the sample without MgATP remains inactive. To assess lipid translocation, outward-facing NBD-lipids are selectively quenched by dithionite, converting them into non-fluorescent ABD-lipid (7-amino-2-1,3-benzoxadiazol-lipid). As a result, the control sample (-ATP) retains higher fluorescence compared to the Mg-ATP sample. Finally, the addition of Triton X-100 (TX-100) solubilizes the proteoliposomes, allowing complete quenching of all remaining NBD-labeled lipids. **(B)**, Cryo-electron image of ΔNC Drs2-Cdc50 proteoliposomes and their size distribution. Bar, 200 nm. To analyze vesicle size, the diameters of 156 vesicles from 9 representative micrographs recorded at a magnification of 92,000× were manually measured using ImageJ software. **(C)**, Specific ATPase activity of ΔNC Drs2 reconstituted in POPC-liposomes with NBD-PS and with (red) or without (dark grey) PI4P, measured using an enzyme-coupled assay in the presence of MgATP and after addition of BeFx. Bars show the mean ± SD (n = 3-16); significance by one-way ANOVA (***p≤0.001, ns p>0.01) **(D)**, Lipid transport by ΔNC Drs2-Cdc50 and catalytically inactive ΔNC E342Q Drs2-Cdc50 mutant reconstituted in POPC liposomes containing 1 mol% NBD-PS and 5 mol% PI4P. Fluorescence traces (mean of 3 technical replicates) were recorded after 30 min incubation at 30 °C with 5 mM MgATP (solid lines) or without MgATP (dashed lines). Dithionite (∼150 s) and TX-100 (∼500 s) were added to quench outer leaflet and total fluorescence, respectively. **(E)**, Quantification of flipped NBD-PS (fluorescence difference at ∼450 s with vs. without MgATP). Bars show the mean ± SD from 3–10 independent reconstitutions; individual data points are shown. ***p ≤ 0.001 (unpaired t-test). **(F)**, NBD-PS transport in proteoliposomes containing ΔNC Drs2-Cdc50 in the presence or absence of PI4P, FL Drs2-Cdc50 with PI4P, ΔNC E342Q Drs2-Cdc50 with PI4P, or in mock (empty) liposomes containing PI4P. NBD-PS transport was analyzed with a fluorescence-based assay, as illustrated in panel A. **(G)**, Substrate specificity of ΔNC Drs2-Cdc50 for various NBD-lipids in PI4P-containing POPC proteoliposomes. In panels D-F, bars show the mean ± SD from 3–10 independent reconstitutions; individual data points are shown. ***p ≤ 0.001 (unpaired t-test).

We reconstituted either ΔNC or FL Drs2-Cdc50 into 1-palmitoyl-2-oleoyl-PC (POPC) liposomes containing 5 mol% PI4P and 1 mol% NBD-labeled lipid, ensuring minimal competition from endogenous substrates. Detergent-mediated reconstitution yielded high protein incorporation (fig. S3A–C), and cryo-EM analysis confirmed mostly unilamellar vesicles with diameters ranging from 30–200 nm (Fig. 2B). Importantly, reconstituted ΔNC Drs2 retained ATPase activity in the presence of PI4P and NBD-PS (Fig. 2C).

We next performed the dithionite quenching assay using the reconstituted proteoliposomes. In this assay, only protein complexes with outward-facing catalytic domains are activated by Mg-ATP and the lipid substrate is transported from the inner to the outer leaflet of the liposome. Thus, active transport upon incubation with Mg-ATP increases the accessibility of the NBD-lipid to dithionite compared to proteoliposomes incubated without Mg-ATP (Fig. 2A). In proteoliposomes containing ΔNC Drs2-Cdc50, PI4P and NBD-PS, fluorescence quenching increased by 10-15% of total fluorescence after incubation with Mg-ATP compared to the control condition, indicating lipid translocation (Fig. 2D-E). Transport was not observed in FL Drs2-, ΔNC Drs2^E342Q^-, or mock-reconstituted liposomes, nor in the absence of PI4P (Fig. 2F), confirming the requirement for autoinhibition relief, PI4P activation and ATP hydrolysis. We next tested whether Drs2 could transport other phospholipids. Proteoliposomes containing ΔNC Drs2-Cdc50, PI4P, and either NBD-PI, -PG, or -PA showed fluorescence quenching comparable to NBD-PS, indicating their active transport. In contrast, no transport was observed for NBD-PC or -PE (Fig. 2G; fig. S4). Taken together, these results demonstrate that Drs2-Cdc50 actively flips a broad range of anionic phospholipids—PS, PI, PG, and PA—but not neutral lipids, consistent with its ATPase activity profile.

### Drs2 is co-purified with substrate lipids

To test the suitability of the ΔNC Drs2-Cdc50 preparation for structural characterization of lipid specificity, we evaluated the influence of possible co-purified substrates on the protein conformation using single particle cryo-EM. Here, we treated the protein with the phosphate analogue aluminum fluoride (AlFx) prior to grid plunging. In the absence of co-purified substrate lipid, the protein should adopt an E1P conformation, whereas a phospholipid occluded ([PL]-E2Pi) conformation would be obtained in the presence of co-purified substrate. Our initial single particle cryo-EM analysis revealed ΔNC Drs2 in a single [PL]-E2Pi conformation, with a distinct lipid density, consistent with PI, in the substrate binding pocket (fig. S5A, C). This indicated the need for removal of substrate lipids during ΔNC Drs2-Cdc50 purification to explore the structural determinants of lipid specificity by cryo-EM.

To remove substrate lipids, we optimized the purification protocol to include a lipid exchange step. Here, the affinity resin-immobilized Drs2-Cdc50 was washed extensively during purification in a buffer containing mixed PC/DDM micelles (0.5 mg/mL DDM and 65 µM PC). This lipid exchanged preparation will be referred to as ΔNC Drs2^PC^-Cdc50. Cryo-EM analysis of ΔNC Drs2^PC^-Cdc50 inhibited by AlFx, resulted in a map consistent with an E1P conformation, indicating successful removal of substrate lipid (fig. S5B, D). Further, the presence of non-substrate PC during purification highlights an important aspect of substrate selectivity, where we only detect map features for transported lipids in the E2P state.

### Drs2-Cdc50 is captured in E2Pi conformation with all substrates

Using ΔNC Drs2^PC^-Cdc50, we investigated the binding mode of the various substrate lipids within the occluded binding pocket of Drs2-Cdc50 by cryo-EM. This was achieved by incubating the protein with substrate lipids and PI4P in detergent before addition of AlFx prior to grid preparation (fig. S7D-E). The respective cryo-EM maps reveal distinct lipid densities for PS, PG, PI, PA and PE captured in the expected [PL]-E2Pi conformation (Fig. 3), validating PS, PG, PI, PA and PE are substrates of Drs2. For PS, PA and PE, a minor class was obtained in an E1P conformation, indicating either more substrate or longer equilibration is needed to fully saturate the binding pocket (fig. S9C-D and fig. S10B).

**Fig. 3:**
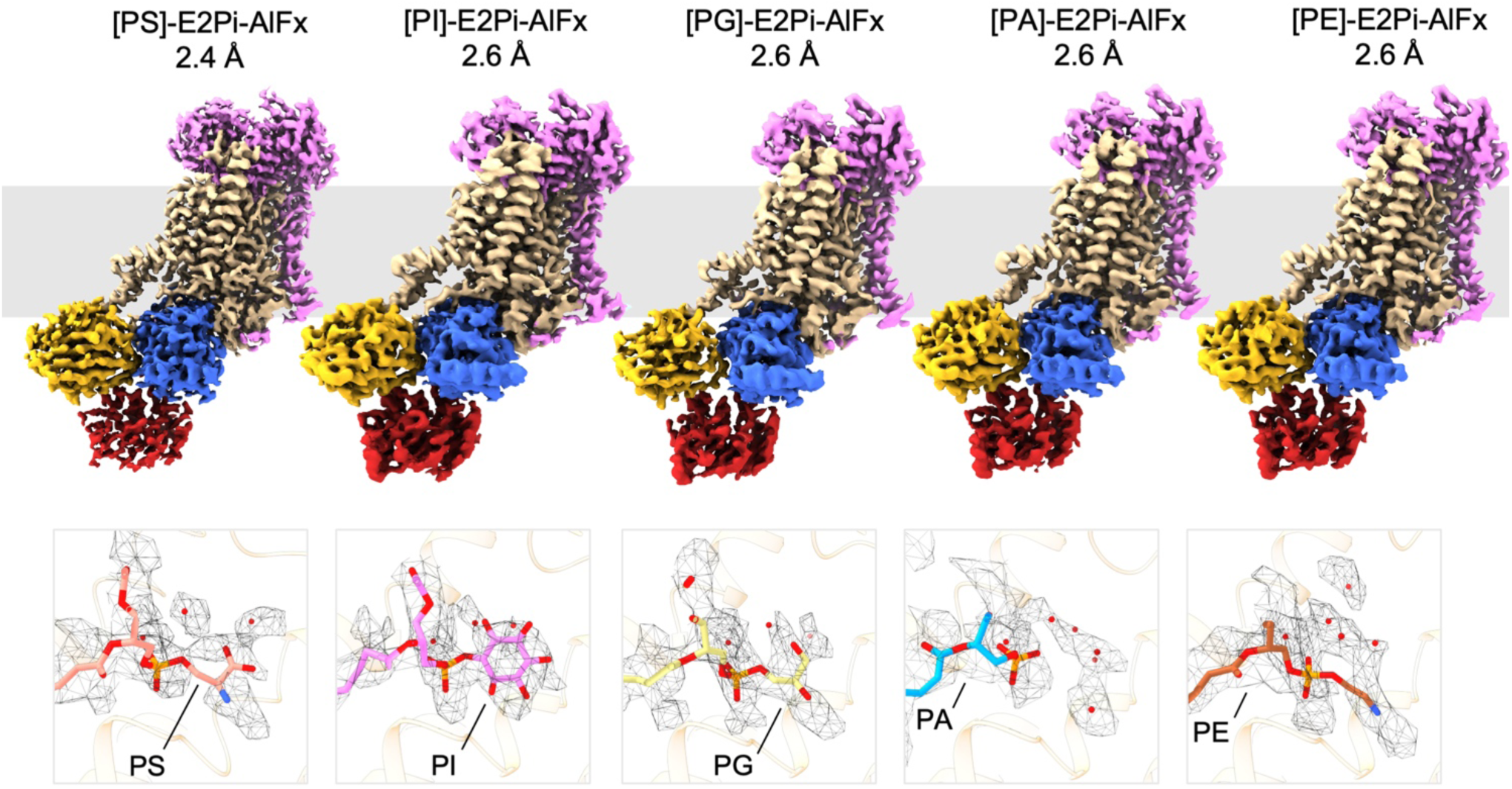
Cryo-EM maps of the Drs2-Cdc50 flippase complex. Top: Cryo-EM maps of Drs2^PC^-Cdc50 in the [phospholipid]-E2Pi-AlFx, lipid occluded state. The maps are colored to highlight key subunits and domains; Cdc50p = pink, TMs = beige, P-domain = blue, A-domain = yellow and N-domain = red. Lipids are colored as follows: PS (salmon), PI (pink), PG (yellow), PA (blue), PE (brown). The identity of the occluded lipid and map resolution is indicated. Inset: model and cryo-EM map densities for substrate lipids and waters in the binding pocket. The contour of the map vs contour used for visualization of the lipid and water densities: PS (6.17 vs 3.3), PI (7.0 vs 6.5), PG (7.4 vs 4.5), PA (6.2 vs 2.8), PE (6.9 vs 3.1).

A key feature in phospholipid recognition is the conserved binding of the phosphate moiety and the *sn*-1 ester carbonyl. Thr249 makes an interaction with the *sn*-1 ester carbonyl of all substrates (fig. S6A-E). The phosphate moiety of the substrate headgroup is coordinated by Asn1050, Ser509 and Asn504. Surprisingly, a distinct density for a water molecule bridging Tyr1023 of TM5 and the phosphate moiety was found in all lipid-occluded maps (Fig. 4 and fig. S6A-E).

**Fig. 4:**
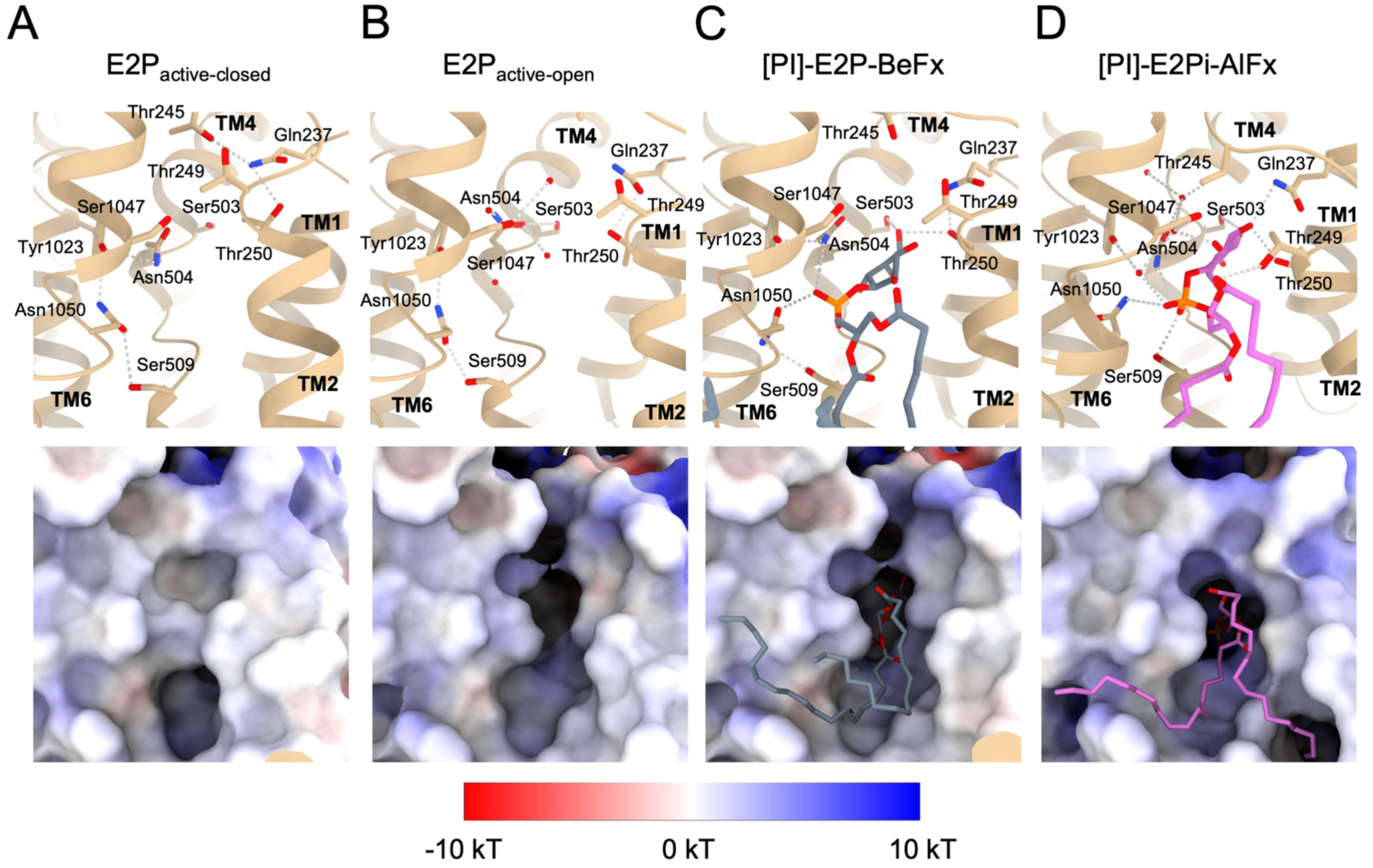
The recognition mechanism. An overview of the binding pocket for **(A)**, E2P_active-closed_ **(B),** E2P_active-open_ **(C)**, [PI]-E2P-BeFx and **(D)**, [PI]-E2P-AlFx, respectively. Top: The lipid-binding site in cartoon representation with residues making contact with PI in the occluded model shown as sticks. The residues in hydrogen bonding distance are connected with a dashed line. The view is from the top of the binding pocket. Bottom: Electrostatic surface showing the closing to opening to occlusion of the substrate binding pocket. The view is into the binding pocket from the back of TM1, 2, 4 and 6.

The lipid headgroups of PS, PI, PG and PE are bound in a similar manner revealing common interacting residues that likely confer specificity (fig. S6A-E and S7A). The serine moiety of PS has direct interactions with residues on TM2 and TM4, namely, Asn504, Ser503 and Thr250, as well as water-bridged interaction with Ser1047, creating a direct interaction interface bridging the static TM4 and TM6 segments, and TM1 and TM2 which move together with the A-domain during lipid occlusion and transport (fig. S6A). The inositol moiety of PI is in hydrogen bonding distance to several residues: Ser1047, Ser503, Thr250 and Gln237 (fig. S6B). The glycerol moiety of PG only makes a direct interaction to Thr250 but interacts with Ser1047 through a water molecule (fig. S6C). PE does not have any interacting residues to the ethanolamine moiety, potentially highlighting why it is a poor substrate of Drs2 (fig. S6D). Together, this points towards Thr249, Thr250, Ser509, Tyr1023 and Asn1050 as being key residues, important for the recognition and occlusion.

On comparison of the various lipids occluded in ΔNC Drs2^PC^ structures, it is evident that the substrate binding pocket is largely invariant in both volume and rotameric position of the amino acid sidechains lining the pocket, irrespective of the occluded substrate (fig. S7B). This may support a role for water molecules in the binding pocket depending on the size of the substrate headgroup (Fig. 3, fig. S8), consistent with the hydrophobic gate model (*53*). This is strongly supported by the PA occluded structure which has the smallest lipid headgroup. Here, we resolve several waters, defining a network of protein-water and water-water interactions from the phosphate group of PA towards Thr245 and Ser1047, emphasizing the potential importance of an ordered water network in lipid occlusion (fig. S6E).

A noted difference is the TM1-TM2 loop, which we resolve in two distinct positions depending on the lipid substrate (fig. S7C). For the lipids where we measure both ATPase activity and transport (PI, PS, PA, and PG), the loop region fully occludes the substrate binding pocket (fig. S7C and S8A-D). For PE, while the TM1 and 2 helical segments are consistent with the other occluded substrates, the TM1-2 loop region is more open, allowing the formation of a continuous channel between the substrate binding site and the luminal compartment (fig S8E). This disconnect between the PE headgroup and the occlusion by the TM1-2 loop may be a crucial element in PE being a poor transport substrate for Drs2.

### The mechanism of substrate recognition

To explore the role of a potential entry gate filter in substate recognition, we examined an earlier step in the transport mechanism by using the phosphate analogue BeFx and treating the ΔNC Drs2^PC^-Cdc50 with and without substrate (fig. S11). Cryo-EM analysis of ΔNC Drs2^PC^-Cdc50 treated only with BeFx, reveals two related conformations differing primarily in their lipid binding site accessibility (fig. S10C). The first has a closed binding pocket (E2P_active-closed_) (Fig. 4A), while the second is consistent with the previously reported E2P_active_ conformation (*12*) where the luminal lipid binding pocket between TM2, TM4 and TM6 is open (E2P_active-open_, Fig. 4B). In the E2P_active-closed_ state, the luminal end of TM2 packs against TM6 to seal the pocket. The transition to the E2P_active-open_ state requires breaking key interactions, including the Thr245-Gln237 and Asn504-Tyr1023 hydrogen bonds, and allows Asn504 to move closer to Ser503, priming the pocket for lipid entry. Interestingly, while ΔNC Drs2^PC^-Cdc50 is prepared in the presence of PC, no additional map density is observed for a phospholipid in the lipid binding site, implying non-substrate lipids have a low affinity to Drs2 or are excluded from entering the binding pocket.

Cryo-EM data of Drs2^PC^-Cdc50 complexed with PI inhibited by BeFx, reveals several conformations including PI-bound E2P and PI-free E2P_active-open_ and E2P_active-closed_ conformations (Fig. 4C and fig. S10D). In the E2P PI-bound state, the binding pocket is open towards the luminal leaflet and is occupied by distinct density for the PI lipid (fig. S11). The phosphate moiety is positioned similar to the fully occluded PI state and is coordinated by Asn504 and Asn1050, while the headgroup is coordinated by Ser503 and Thr250 (Fig. 4C). For lipid occlusion, these interactions likely trigger a rearrangement of the internal hydrogen-bonding network, involving Gln237, Ser509, and Tyr1023 and ordered water molecules (Fig. 4C-D, fig. S11C). Based on these structures, we propose a two-step mechanism where initial lipid recognition and binding is via residues on the largely static TM4 and TM6 which interact with the phospholipid headgroup. Rearrangement of TM1-2 provides additional headgroup interacting residues (Thr249 and Gln237) that subsequently leads to occlusion in a hydrated cavity (fig. S8B and S11).

## Discussion

Assessing the transport specificity of lipid flippases has remained challenging to date, partly due to the difficulty of monitoring lipid transport *in vitro*. Using the reconstitution procedure outlined here, we successfully incorporated functional Drs2-Cdc50 in unilamellar vesicles suitable for lipid transport experiments. Notably, our reconstitution protocol differs in key aspects from previously published methods for functional reconstitution of Drs2-Cdc50, such as lipid composition of the proteoliposomes, or the type/concentration of detergent used for detergent-mediated reconstitution (*36*, *54*). Reconstituted Drs2 exhibits ATPase activity towards NBD-labeled phospholipid analogues, which is particularly important for establishing a lipid transport assay that relies on the transport of fluorescently labelled phospholipids. While previous attempts to measure flippase-mediated lipid transport using a similar assay often detected fluorescence changes of a few percent (*36*, *54*), we reliably observed changes up to 15%, demonstrating high sensitivity and robustness of our assay.

This improved sensitivity enabled us to study the transport of various lipids and show that Drs2-Cdc50 transports not only its known substrate PS, but also PI, PG and PA. However, while our cell-based transport assays indicated that Drs2-Cdc50 facilitated transbilayer movement of NBD-PE, we failed to detect such transport *in vitro*. The fundamental reason for this apparent discrepancy is unclear. However, our cryo-EM derived structures of Drs2 in the presence of PE yielded a partly occluded lipid bound structure, where the TM1-2 loop is not fully closed over the luminal opening resulting in a channel continuous with the luminal milieu (fig. S8E), indicating PE may be a potential albeit poor transport substrate. Structural studies in the presence of the non-substrate PC resulted in an AlFx stabilized E1P state (fig. S11A). We propose that the final positioning of this loop region relies on a stable binding of the lipid headgroup. The headgroups of PS, PI and PG allow for multiple interactions to define binding and limit movement, while PE can likely sample multiple conformations in the lipid binding site. We speculate that only one of these conformations could support full occlusion by the TM1-2 loop and transport of the lipid. Additionally, we note that previous studies with TGN membranes harboring temperature-sensitive alleles of *DRS2* also did not reveal any significant role of Drs2 in NBD-PE transport (*34*). Furthermore, the quadruple yeast mutant used in our *in vivo* transport assays still contains Neo1, which has been shown to promote PS and PE asymmetry at the plasma membrane (*43*). Due to the dual role of P4-ATPases in lipid transport and vesicle formation in the secretory pathway, we cannot rule out that Neo1-dependent transport of PE at the PM is upregulated in yeast P4-ATPase mutants. An alternative explanation could stem from the experimental conditions used in our *in vitro* measurements. While our ATPase and transport assays were conducted at pH 7, transport of PE by mammalian ATP8A2 appeared to be largely diminished when the pH was lowered from 7.5 to 6.7, suggesting that PE transport is dependent on the ionization of the headgroup (*55*).

The unexpectedly broad specificity of Drs2-Cdc50, a model flippase long considered highly specific for PS, strongly suggests that the substrate profiles of other P4-ATPases warrant careful re-examination. The specificity of many flippases has been assigned based primarily on cell-based assays using fluorescent lipid analogs. As our own results with PE and PA demonstrate, this approach can be confounded by the complex cellular environment, including the compensatory activity of other endogenous transporters and trafficking perturbations regulatory networks, leading to potentially incomplete conclusions. The clear discrepancy we observed between 1) the apparent PE transport in yeast mutants and the lack of activity in our purified, reconstituted system and 2) the lack of in-cell NBD-PA uptake and low stimulation of ATPase activity but robust transport in proteoliposomes underscores the necessity of *in vitro* validation to assign substrate specificity definitively. This paradigm is further supported by recent revisions to the substrate profile of other flippases, such as ATP8B1 and ATP10A, which have also been found to transport PI (*26*). Therefore, a multi-pronged strategy, combining in-cell observations with robust *in vitro* transport assays and structural analysis, is essential to build an accurate understanding of the physiological roles of this critical family of lipid transporters.

Our studies reveal that co-purification of transport substrates and the presence of regulatory features provide a significant challenge in the structural and functional studies of lipid flippases, where structures of lipid occluded or off-cycle transport states have the potential to be incorrectly interpreted. Furthermore, the presence of co-purified substrate lipids provides basal ATPase activity and the potential to confound *in vitro* assays exploring lipid specificity. By utilizing a lipid exchange protocol to a non-substrate lipid coupled with the removal of auto-regulatory segments, we provide a robust strategy to explore the breadth of lipid specificity and transport by lipid flippases.

Structural studies of the lipid-exchanged Drs2 have provided new understanding in the mechanism of lipid recognition and transport. The invariant lipid binding pocket precludes an induced fit mechanism for substrate recognition. Instead, it points to a requirement of a hydrated lipid model for transport. The flippase recognizes not the lipid in isolation, but an entity formed by its polar headgroup and a structured solvation shell defined by the flippase and transport lipid. The transported substrates (PS, PI, and PG) form a stable hydrated ligand that triggers occlusion. Remarkably, PA achieves this by organizing a structural water network that functionally mimics a hydrated lipid headgroup. Initial substrate recognition and binding is via residues in the largely rigid TM4 and 6 (Asn504, Ser509, Tyr1023 and Asn1050), of which Ser509 and Asn1050 are conserved (figure S16). Occlusion and transport then occur via movements in the structural elements (TM1 and 2) coupled to the A-domain. The observed substrate binding mode is similar to the asymmetric substrate coordination found in the major facilitator superfamily of secondary active transporters (*56*, *57*).

The broad specificity of Drs2 is supported by the invariant cavity that houses the lipid headgroup during transport, which can only occlude when discrete interactions between the substrate lipid headgroup, solvating waters and the various structural elements of Drs2 can be satisfied. On comparison to ATP8A1 (PDB 6K7M), ATP8B1 (PDB 8OXA and 8OXB) and ATP11C (PDB 7BSV), we observe a similar positioning of PS in the binding pocket (fig. S12A and fig. S16). The phosphate moieties are aligned, and they are coordinated by the same set of residues, namely the conserved TM6 asparagine and the serine from the PISL motif (fig. S12B-E). Asn504 coordinates the phosphate moiety in Drs2, and this residue, and the corresponding interaction, are only conserved in ATP8A1. In ATP8B1, this locus is a threonine, while it is a phenylalanine in ATP11C. In ATP11C, this phenylalanine, together with a more closed luminal end of TM2, results in a narrower cavity (fig. S13E) which may favor substrates with smaller headgroups, e.g., PS and PE. In the TM1-TM2 loop, ATP8B1 has a leucine instead of the Thr245, providing a less polar binding pocket. Together with the Thr398 on TM4, these residues would help create an environment for PC to bind. Here, the corresponding Asn504 on Drs2 would sterically clash with a PC, if positioned as in ATP8B1 (PDB 8OXB) (fig S12F).

The findings presented here reveal that Drs2-Cdc50 transports PI from the luminal to the cytosolic leaflet of yeast TGN membranes. Although it has been demonstrated that PI is predominantly confined to the cytosolic leaflet of the PM in erythrocytes, studies on the transbilayer asymmetry of PI in endomembranes are scarce. However, a recent report indicates that PI is strongly shifted toward the cytosolic leaflet in the TGN secretory vesicles of live HeLa cells (*58*). The transbilayer distribution of PI in yeast remains unknown, though we speculate that Drs2-Cdc50 may play a critical role in this process. Importantly, PI is the precursor of PIPs, including PI4P, which is primarily located to the TGN, and to a lesser extent to the PM (*59*). The subcellular distribution of PIPs is maintained through a tight spatiotemporal control of their synthesis and degradation by PI kinases and phosphatases, respectively. The type III PI 4-kinase Pik1 is responsible for Golgi synthesis in the yeast TGN, while Sac1, a PI4P phosphatase cycles between the ER and the early Golgi (*60*). Owing to the various cellular roles of PIPs, particularly PI4P, in cell signaling, membrane trafficking and control of membrane protein activity (*61*), the ability of Drs2-Cdc50 to transport its precursor PI is of special interest. In agreement with a central role of Drs2-Cdc50 in PIP metabolism, a synthetic genetic interaction between *drs2Δ* and *pik1Δ* has previously been observed (*62*). Moreover, *in vivo* cross-linking coupled with mass spectrometry uncovered a physical interaction between Drs2 and PIP-metabolizing enzymes, including Sac1, the *myo*-inositol transporter Itr1 and the *myo*-inositol 1-phosphate synthase Ino1 (*63*). The formation of TGN-derived secretory vesicles en route to the plasma membrane, as well as AP1-mediated transport from the TGN to the endosomes, is dependent on PI4P (*64*, *65*). Similar to PI4P, Drs2-Cdc50 contributes to budding of exocytic vesicles from the TGN (*38*) and is required for bidirectional transport between the TGN and the early endosomes, as well as for AP-1 function in this pathway (*37*, *66*). This raises the question as to whether the trafficking phenotypes observed upon Drs2 inactivation are not associated with reduced PI4P synthesis in the cytoplasmic leaflet of the TGN, a consequence of defective PI transport by Drs2-Cdc50. Given that the P4-ATPases ATP8B1 and ATP8B2 have recently been suggested to transport PI, this hypothesis aligns with findings indicating a reduction in PI(4,5)P_2_ levels in the PM of *ATP8B1/2* knockout cells when stimulated by Gq-coupled receptors, compared to parental HeLa cells (*26*).

In summary, this study sheds light on the unexpectedly broad specificity of the Drs2-Cdc50 complex, the ortholog of mammalian ATP8A1-CDC50 complex, for anionic lipids. Using complementary *in vivo* and *in vitro* approaches, the results clearly show that Drs2-Cdc50 transports PS, PI, PG and PA. Such Drs2-catalyzed active transport may have significant impact on the transbilayer distribution of these anionic lipids in the yeast TGN and in turn, on PI4P homeostasis. The present findings also highlight the need to carefully reevaluate the substrate specificity of other P4-ATPase lipid flippases.

## Materials and Methods

### Lipids and detergents

POPC (1-palmitoyl-2-oleoyl-*sn*-glycero-3-phosphocholine), POPS (1-palmitoyl-2-oleoyl-*sn*-glycero-3-phospho-L-serine), POPI (1-palmitoyl-2-oleoyl-*sn*-glycero-3-phosphoinositol), POPG (1-palmitoyl-2-oleoyl-*sn*-glycero-3-phospho-{1’-rac-glycerol}), POPA (1-palmitoyl-2-oleoyl-*sn*-glycero-3-phosphate), POPE (1-palmitoyl-2-oleoyl-*sn*-glycero-3-phosphoethanolamine), DOPE (1,2-dioleoyl-*sn*-glycero-3-phosphoethanolamine), DOPS (1,2-dioleoyl-*sn*-glycero-3-phospho-L-serine), DOPA (1,2-dioleoyl-*sn*-glycero-3-phosphate), DOPG (1,2-dioleoyl-*sn*-glycero-3-phosphoglycerol), 18:1-12:0 NBD-PS (1-oleoyl-2-{12-[(7-nitro-2-1,3-benzoxadiazol-4-yl)amino]dodecanoyl}-*sn*-glycero-3-phosphoserine), 16:0-12:0 NBD-PS (1-palmitoyl-2-{12-[(7-nitro-2-1,3-benzoxadiazol-4-yl)amino]dodecanoyl}-*sn*-glycero-3-phosphoserine), 18:1-12:0 NBD-PG (1-oleoyl-2-{12-[(7-nitro-2-1,3-benzoxadiazol-4-yl)amino]dodecanoyl}-*sn*-glycero-3-[phospho-rac-(1-glycerol)]), 18:1-12:0 NBD-PA (1-oleoyl-2-{12-[(7-nitro-2-1,3-benzoxadiazol-4-yl)amino]dodecanoyl}-*sn*-glycero-3-phosphate), 18:1-12:0 NBD-PE (1-oleoyl-2-{12-[(7-nitro-2-1,3-benzoxadiazol-4-yl)amino]dodecanoyl}-*sn*-glycero-3-phosphoethanolamine), 16:0-12:0 NBD-PC (1-palmitoyl-2-{12-[(7-nitro-2-1,3-benzoxadiazol-4-yl)amino]dodecanoyl}-*sn*-glycero-3-phosphocholine), and brain PI(4)P (L-α-phosphatidylinositol-4-phosphate) were purchased from Avanti Polar Lipids. 16:0-12:0 NBD-PI was synthesized in-house by transphosphatidylation of 16:0-12:0 NBD-PC with *myo*-inositol, using a microbial phospholipase D variant with positional selectivity for the 1-OH group of *myo*-inositol, as previously described (*67*). Triton X-100 (TX-100) was from Sigma-Aldrich and n-Dodecyl-β-D-Maltoside (DDM) was from Anatrace.

### DNA constructs

Full-length Drs2 was N-terminally tagged with a tobacco etch virus (TEV)-cleavable biotin acceptor domain (BAD). The ΔNC Drs2 variant was C-terminally tagged with a thrombin-cleavable BAD domain. The sequence included an engineered thrombin cleavage site inserted after residue 1247 in the C-terminal tail, in addition to the natural thrombin cleavage sites at residues 104 and 1290, allowing for the removal of Drs2 N- and C-terminal extensions by enzymatic cleavage with thrombin. The catalytically inactive mutant ΔNC E342Q was generated from the ΔNC Drs2 construct by site-directed mutagenesis. For *in vivo* lipid uptake assays, untagged versions of full-length Drs2 and its beta subunit Cdc50 were inserted by restriction-based cloning on each side of the bi-directional GAL1-10 promoter of plasmid pRS423-GAL (*68*). Functional complementation assays used a modified version of pRS423-GAL bearing the *DRS2* gene (*46*).

### Lipid uptake assays in living yeast cells

NBD-lipid transport was measured as previously described (*44*). Briefly, yeast strain ZHY709 (*MATa his3 leu2 ura3 met15 dnf1Δ dnf2Δ drs2*∷LEU2) (*69*) was transformed using the lithium acetate/single-stranded carrier DNA/polyethylene glycol method (*70*). Transformants were selected onto SCD-His (0.7% yeast nitrogen base without aminoacids, 2% glucose, 1x yeast synthetic drop-out medium supplements without histidine (Sigma, cat. no. Y1751), 2% agar). Drs2 expression was induced in liquid SCG-H cultures containing 2% galactose instead of glucose. Cells were collected at OD_600_=0.3-0.5 an concentrated in SCG-H medium to OD_600_=10. Cells were divided into 125 µL aliquots and latrunculin A was added at a final concentration of 16 µM. After a 5-min incubation at 30℃, NBD-lipids were added at a final concentration of 24 µM. Cells were then incubated 30℃ for 30 minutes, transferred to and ice-water bath, and immediately washed 3 times with SSA+BSA (same as SCG-H but with 2% sorbitol instead of galactose and containing 3% bovine serum albumin and 13 mg/mL sodium azyde) to eliminate excess lipids. Fluorescence intensity was measured using a FACScalibur flow cytometer (BD Biosciences) equipped with a 488 nm laser. Propidium iodide was used as a live/dead staining and at least 20,000 living cells were counted for each yeast population. Data analysis was carried out using FlowJo software (FlowJo LLC).

### Functional complementation assays in yeast

Yeast strain TPY127 (*MAT*α *sec6-4* TPI1::SUC2::TRP1 *ura3-52 his3-Δ200 leu2-3-112 trp1-1Δdrs2-41::loxP Δdnf1::loxP Δdnf2::loxP Δdnf3::loxP-HIS3-loxP*) (*42*) was transformed and selected as above. Fresh transformants (5-10 colonies) were incubated for 6 hours at 30℃ with150 rpm shaking in 2 mL of SCGF-H medium (same as SCG-H but with addition of 2% fructose) before measuring OD_600_. Serial 1:10 dilutions, starting at OD_600_ = 0.1, were dropped (5 µL) on SCD-H and SCGF-H plates, as controls, and on SCGF-H plates supplemented with 0.1 µg/mL papuamide A, 1.5 µM duramycin, or 2.5 µg/mL miltefosine, respectively. Plates were incubated at 28℃ for 4-5 days.

### Protein expression and purification

BAD-tagged Drs2 was overexpressed with either N- or C-terminally His-tagged Cdc50 using a single plasmid (pYeDP60) for co-expression of both proteins in the yeast *Saccharomyces cerevisiae* strain W303.1b/*GAL4* (*MATa*, *leu2*, *his3*, *trp1*::*TRP1-GAL10-GAL4*, *ura3*, *ade2-1*, *canr*, *cir+*). Yeast transformation with the respective plasmids was performed using the lithium acetate/single-stranded carrier DNA polyethylene glycol method (*70*). Transformed cells were grown on SD-Ura medium (2% w/v dextrose, 0.17% w/v yeast nitrogen base, 0.5% w/v (NH_4_)_2_SO_4_, 55 µg/mL adenine SO_4_, 55 µg/mL L-tyrosine, 20 µg/mL L-arginine, 10 µg/mL L-histidine, 60 µg/mL L-isoleucine, 60 µg/mL L-leucine, 40 µg/mL L-lysine, 10 µg/mL L-methionine, 60 µg/mL L-phenylalanine, 50 µg/mL L-threonine, 40 µg/mL L-tryptophan) selective plates (2% w/v agar) for 3–5 days at 28 °C. Single colonies were inoculated into 5 mL SD-Ura and grown for 24 h at 28 °C, 180 rpm. The pre-culture was used to inoculate 50 mL SD-Ura (starting OD₆₀₀ of 0.1), followed by incubation under the same conditions. Cultures were then transferred to rich medium (2% w/v yeast extract, 2% w/v bactopeptone, 1% w/v dextrose, 2.7% v/v ethanol) at an OD₆₀₀ of 0.05, and incubated for 36 h at 28 °C, 130 rpm. After cooling on ice for 10-15 min, protein expression was induced by adding 2% w/v galactose. Cells were incubated for 13 h at 18 °C, 130 rpm, then supplemented with another 2% w/v galactose and incubated for an additional 5 h under the same conditions. Cells were harvested by centrifugation (1,000 g, 10 min, 4°C) and washed with ice-cold water and buffer (50 mM Tris-HCl pH 7.5, 1 mM EDTA, 100 mM KCl, 0.6 M sorbitol). Unless otherwise specified, all subsequent steps were performed on ice or at 4°C. Cell pellets were resuspended at a 1:1 w/v ratio in lysis buffer (50 mM Tris-HCl pH 7.5, 1 mM EDTA, 0.6 M sorbitol, 2 mM PMSF, and protease inhibitors) and lysed with 0.5 mm glass beads using a planetary mill (Pulverisette 6, Fritsch). The lysate was cleared by low-speed centrifugation (1,000 g, 20 min), and the supernatant S1 was subjected to medium-speed centrifugation (20,000 g, 20 min), resulting in supernatant S2 and pellet P2. S2 was ultracentrifuged (125,000 g, 1 h), and the obtained membrane pellet P3 was resuspended in 20 mM Hepes-Tris pH 7.4, 0.3 M sucrose, 0.1 mM CaCl_2_. The total protein concentration in P3 was determined with a bicinchoninic acid assay, and expression of Drs2 and Cdc50 was confirmed by western blotting.

To purify WT Drs2, ΔNC Drs2, and ΔNC Drs2^E342Q^, P3 membranes were diluted to 5 mg/mL total protein in SSR buffer (50 mM MOPS-Tris pH 7, 100 mM KCl, 5 mM MgCl_2_) supplemented with 20 % w/v glycerol, 1 mM PMSF and protease inhibitors (either 1x EDTA-free protease inhibitor cocktail, Sigma-Aldrich, or 1 µg/mL each of pepstatin, leupeptin and chymostatin). Membranes were solubilized with 5 mg/mL DDM for 15 min under stirring and centrifuged (125,000 g, 1 h) to remove insoluble material. The supernatant was incubated with streptavidin sepharose resin (0.33 ml per mg Drs2) for 2 h at 4 °C with slow stirring. The resin was washed 3-4 times with 3 column volumes (CV) SSR buffer supplemented with 20% w/v glycerol and 0.5 mg/mL DDM. For elution, Drs2-Cdc50 complexes were cleaved overnight on resin using TEV protease (60 µg/mL resin) for full-length Drs2, or thrombin (4 U/mL) for ΔNC variants. Eluates were pooled and concentrated (100 kDa MWCO). For ΔNC Drs2 and ΔNC Drs2^E342Q^, a second thrombin incubation (80 U/mL) was performed in the presence of 50 µg/mL POPS and 25 µg/mL PI4P for 2 h at 20 °C, to cleave the N- and C-terminal extensions. This resulted in ΔNC Drs2 comprised of residues 104-1247. Protease activity was stopped with 2 mM PMSF. For size exclusion chromatography (SEC), samples were loaded on a Superdex 200 Increase 10/300 GL column equilibrated with SSR buffer supplemented with 0.5 mg/mL DDM and 1 mM DTT. Protein-containing fractions were pooled, frozen in liquid N_2_, and stored at -80 °C. Protein concentrations were determined by measuring absorbance at 280 nm using a spectrophotometer (NanoDrop) and extinction coefficients of *ε*=237 mM^-1^.cm^-1^ for FL Drs2-Cdc50 and *ε*=225 mM^-1^.cm^-1^ for ΔNC Drs2-Cdc50 and ΔNC Drs2^E342Q^- Cdc50. Extinction coefficients were calculated based on the amino acid sequence.

### Liposome preparation

Powdered POPC was pre-cleaned prior to liposome preparation. The lipid was first dissolved in chloroform (∼2.5 mL/mg) in a balloon-shaped glass tube. The solution was dried under vacuum using a rotary evaporator (Rotavapor R-200, BÜCHI) until a uniform, thin lipid film formed on the glass surface. This dried film was then hydrated in Milli-Q water (2–3 mL/mg lipid), vortexed, sonicated in a water bath to fully resuspend the lipids, and frozen in liquid nitrogen. The suspension was lyophilized overnight to yield a dry, chloroform-free lipid powder. For liposome formation, the resulting lipid powder was resuspended by vortexing in SSR buffer to a final lipid concentration of 5 mM, aliquoted and stored at -20 °C until use. Before use, each batch was solubilized with Triton X-100, and the detergent-to-lipid ratio required to reach the onset of full solubilization (typically 2.5–3 w/w) was determined by measuring absorbance at 500 nm.

### Drs2 reconstitution in proteoliposomes

SEC-purified Drs2-Cdc50 was reconstituted into POPC liposomes at room temperature. Liposomes (5 mM lipids) were solubilized with Triton X-100 (TX-100) at a 2.5-3.0 (w/w) detergent-to-lipid ratio and stirred gently for 30 min. Separately, C12-NBD-labeled phospholipids (1 mol% final) were dried under vacuum or argon stream. Solubilized liposomes were added to the dried NBD-lipids along with protein (0.1 mg/mL final, ∼500 nM) and SSR buffer to adjust the lipid concentration to 2.5 mM, yielding a protein/lipid molar ratio of 1:5000, assuming no loss of proteins or lipids during the reconstitution.

If required, PI4P (5 mol%) was included by drying it with the NBD-lipids or adding from a PI4P/DDM stock. The protein-lipid-detergent mixture was gently stirred for 30 min to ensure complete solubilization of the NBD-lipids. Detergent was removed with Biobeads SM-2 (Bio-Rad), which were pre-washed with methanol and water, and stored in SSR buffer at 4 °C until use. Detergent removal was performed at room temperature, in three steps. First and second, Biobeads were added at a Biobead/detergent w/w ratio of 10:1 followed by incubation for 2 h with gentle stirring. Third, the amount of Biobeads was doubled (Biobead/detergent w/w ratio of 20:1) followed by incubation for 1 h with gentle stirring. Empty liposomes (mock) were prepared similarly, but an equal amount of SSR buffer was added instead of protein. Proteoliposomes and empty liposomes were stored at 4 °C and used within 3-4 days.

### Cryo-electron microscopy of proteoliposomes

4 μL of solution of proteoliposomes diluted at 0.25 mg/mL in SSR buffer were deposited on glow-discharged carbon-Formvar lacey grids (Ted Pella, USA), blotted from the back side, and flash frozen in liquid ethane with an EM-GP2 Leica plunger at 90% humidity. Cryo-EM images were acquired with a Glacios cryo-electron microscope (Thermo Fisher, USA) operating at 200 kV with a Falcon IVi camera and in low dose mode.

### Flotation assay

Proteoliposomes and mock liposomes were subjected to flotation assays to assess the colocalization of Drs2-Cdc50 and NBD-lipids, indicating successful incorporation of proteins in vesicles. Sucrose was dissolved in SSR buffer, and gradients were prepared in polycarbonate ultracentrifuge tubes as follows: 200 µL sample was overlaid with 200 µL 60% w/v sucrose, 300 µL 24% w/v sucrose and 200 µL SSR buffer. Gradients were centrifuged at 55,000 rpm for 90 min at 4 °C (TLS 55 rotor, Beckmann). After ultracentrifugation, three fractions (B – bottom, M – middle, T – top) were collected from the bottom of the gradient with a Hamilton syringe. NBD fluorescence of each fraction was measured (1:20 dilution in SSR buffer) using a fluorescence microplate reader (GENios, TECAN, 485 nm excitation, 535 nm emission). Background fluorescence from SSR buffer was subtracted. Measurements were performed in triplicate. The protein content in each fraction was quantified by SDS-PAGE: 40 µL of each fraction was mixed with 10 µL 5x SDS loading buffer, and 35 µL of this mixture was loaded on a 10% acrylamide SDS-gel.

### ATPase activity measurements

ATPase activity of detergent-solubilized and SEC-purified ΔNC Drs2-Cdc50 was measured using a colorimetric assay based on the formation of a phosphomolybdate complex, which absorbs at 860 nm at low pH. Reactions were performed in SSR buffer in a total volume of 120 µL with 10 µg/mL ΔNC Drs2-Cdc50, 1 mg/mL DDM, 25 µg/mL PI4P, and varying concentrations of lipids (lipid/detergent molar ratios up to 0.3) added as a solubilized lipid/detergent mix (10 mg lipids per 50 mg DDM). Reactions were initiated by adding 2 mM MgATP and incubated at 30 °C for 10 min. Subsequently, 33 µL of the reaction was mixed with 17 µL of 10% SDS in 96-well plates, followed by the addition of 200 µL of colorimetric reagent (3 mM zinc acetate, 7 mM ammonium molybdate in 10% sodium ascorbate, at pH 5). After 30 min incubation at 20 °C, absorbance was measured at 860 nm. Liberated phosphate was quantified using a K₂HPO₄ standard curve (0–0.4 mM). The rate of ATP hydrolysis was constant for at least 20 min under the assay conditions (data not shown). After subtracting background activity (measured in the absence of lipids), the corrected ATP hydrolysis rates were plotted against lipid/detergent molar ratios and data points were fitted to a Michaelis-Menten-equation (Y = Vmax*X/(Km + X)) using GraphPad Prism, yielding V_max_ and K_m_ values.

ATPase activity of SEC-purified or reconstituted Drs2-Cdc50 was measured using an enzyme-coupled assay in which ATP hydrolysis is coupled to NADH oxidation in the presence of lactate dehydrogenase (LDH), pyruvate kinase (PK) and phosphoenolpyruvate (PEP). For SEC-purified Drs2-Cdc50, ∼20 µg protein was added to 2 mL assay medium (0.04 mg/mL PK, 0.1 mg/mL LDH, 1 mM MgATP, 1 mM PEP, 1 mM NaN_3_, 1 mM DTT and 0.3 mM NADH in SSR buffer) supplemented with 0.5 mg/mL DDM, 50 µg/mL POPS and 25 µg/mL PI4P. Measurements were performed at 30 °C. NADH absorbance at 340 nm was continuously monitored and the rate of absorbance decrease was converted into specific ATPase activity as described (*31*). In the case of FL Drs2, 10 µg/mL trypsin was added directly to the cuvette to reveal full activity. 0.5 mM BeFx was finally added to inhibit ATPase activity.

For reconstituted Drs2-Cdc50 in proteoliposomes, 40 µL sample was mixed with 2 mL assay medium and NADH absorbance was monitored as before. Drs2 concentration in proteoliposomes was estimated by density analysis on SDS-PAGE or western blot and compared to a Drs2 standard curve.

### Lipid transport assays in proteoliposomes

NBD-lipid transport in proteoliposomes was measured by dithionite reduction of NBD fluorescence. Liposomes (30 µL) were incubated with or without 5 mM MgATP at 30 °C for 30 min. After incubation, 20 µL of the sample was immediately transferred into 2 mL SSR buffer in a temperature-controlled quartz cuvette (20 °C, constant stirring). NBD-fluorescence was recorded at 530 nm (excitation 470 nm, 1 s integration) using a Horiba Fluorolog fluorometer. Bandwidths were set to 2-4 nm, so that the initial signal is between 1-2x10^6^ counts per second. After around 150 s, 40 µL of freshly prepared 1 M dithionite (in 0.5 M Tris) was added to yield a final dithionite concentration of 20 mM, and fluorescence quenching was followed for ∼350 s. After ∼500 s, 0.3 mg/mL TX-100 was added to fully solubilize liposomes, resulting in complete fluorescence loss. Fluorescence intensity was normalized according to *F_norm_= (F-F_0_)*100/(F_init_-F_0_)*, where *F* is the fluorescence at each time point, *F_init* is the average fluorescence over 12 s before dithionite addition, and *F₀* is the average over ∼60 s after TX-100 addition. The percentage of NBD-phospholipid transported from the inner to the outer proteoliposome membrane was calculated as the difference in *F_norm_* after 450 s between samples incubated with and without MgATP.

### Purification and lipid exchange of ΔNC Drs2-Cdc50

The membranes were diluted to 5 mg/mL of total protein in SSR buffer (50 mM MOPS-Tris pH 7, 100 mM KCl and 5 mM MgCl2, 1 mM DTT and 20% (w/v) glycerol) containing 1 mM PMSF and 1 μg/mL of leupeptin, pepstatin and chymostatin. They were solubilized for 15 min and centrifugated at 42,000 rpm for 1h (Sorvall WX+ ultracentrifuge, T647.5 rotor). The solubilized membranes were batch-bound to streptavidin resin for 1 hour at 4◦C and washed with 2 column volumes (CV) of SSR containing 0.5 mg/mL DDM, 1 mM PMSF and 1 μg/mL of leupeptin, pepstatin and chymostatin. The resin was washed in batch with ten times with 5 CV of SSR containing 0.5 mg/mL DDM with 65 µM POPC, with a 10 min incubation at RT. The resin was washed with 2 CV SSR with 0.2 mg/mL Lauryl Maltose Neopentyl Glycol (LMNG) and subsequently washed with 10 CV SSR with 0.1 mg/mL LMNG. The protein was cleaved from the resin overnight (ON) with 4 U/mL thrombin at 4◦C. The protein was eluted from the resin in 5 CV SSR with 0.1 mg/mL LMNG and then concentrated 40-fold in a 100 kDa MWCO spin column (VivaSpin). To generate the truncated ΔNC Drs2-Cdc50, the autoregulatory domain was removed by incubating with 25 μg/mL phosphatidylinositol-4-phosphate (PI4P) and 5 U/mL thrombin for 1 hour at RT. SEC was performed using a Superdex 200 increase 10/300 (Cytiva Life Sciences) on an Äkta Go (Cytiva Life Sciences) using SSR with 0.03 mg/mL LMNG. Peak fractions were analyzed by SDS-PAGE with appropriate fractions pooled, resulting in a protein concentration of 1 mg/mL in 600 µL.

### Grid preparation

UltrAuFoil 1.2/1.3 300 mesh (Quantifoil), were glow-discharged on a GloQube Plus Glow Discharge for 45 sec at 15 mA. Before addition of the sample to the grid, 0.8 mg/mL ΔNC Drs2-Cdc50 in LMNG was incubated with 0.2-0.3 mg/mL phospholipid for 2 hours and treated with 2 mM AlFx or 2 mM BeFx, respectively, for 20 min. The inhibitors were prepared in a 10X stock from a mix of 1:10 KF and AlCl_3_ (AlFx) or 1:10 KF and BeSO_4_ (BeFx). Following, 3 μl of sample was added to the grid and vitrified on a Vitrobot IV (Thermo Fisher) at 4 °C and 100% humidity. The conditions for blotting were blot force 0, blot time 3 or 3.5 sec and wait time of 10 sec.

### Cryo-EM data collection

Cryo-EM data was collected on a Titan Krios with an X-FEG operated at 300 kV equipped with a Gatan K3 camera and a Bioquantum energy filter operated at a slit width of 20 eV. Movies were collected using aberration-free image shift data collection (AFIS) in EPU (Thermo Fisher Scientific) with a calibrated pixel size of 0.647 Å (magnification of 130,000x), with a fluence of 60 electrons per Å^2^. For data collection, a defocus range between -1.8 and -0.8 μm was used.

### Data processing and model building

Cryo-EM data was processed in cryoSPARC (v4) (*71*, *72*) and a general processing outline is presented in fig. S14. Patch Motion Correction and Patch CTF were performed before micrograph curation on the basis of CFT estimations, defocus, total frame motion and ice thickness. Particles were picked by blob picking, from which ab-initio volumes were generated. The volumes were used in Heterogeneous Refinement, and the best volume from Heterogeneous Refinement was used to create 2D templates and re-pick particles by template picking. New ab-initio volumes were generated and used in Heterogeneous Refinement to sort out junk and broken particles. 3D Classification was performed to sort conformational heterogeneity and sub-optimal particles. For 3D Classification, a mask around TM1-TM4 including the A-domain was used. Class similarity was set to 0.8 with force hard classification turned on. The final particle stack was used to perform Non-Uniform Refinement, with dynamic mask start resolution at 1Å. The resulting map and particles were polished using Reference based Motion Correction and subjected to a final non-uniform refinement. Resulting map resolution was determined using the gold standard fourier shell correlation (FSC, 0.143 cutoff). Local map resolution estimates were determined using CryoSPARC. An overview of all maps with local resolution estimates and plots respectively depicting FSC, orientation distribution and anisotropy are presented in fig. S15.

Previous models of Drs2-Cdc50 in E1P (7OH5), E2Pactive (6ROJ) and E2Pi (PDB: 7OH6) were used as starting models and fit into the new respective maps first by Namdinator (*73*), and then by Isolde (*74*), using cryo-EM maps sharpened using PHENIX Autosharpen (*75*). Manual modelling of protein and ligands was perfomed in Coot (v 0.9.8.93) with several iterative rounds of Real-Space refinement as implemented in PHENIX (*76*). Lipids were modelled based on the type of lipid added during sample preparation. For Real-Space refinement, hydrogen addition and ligand restraints were generated by the ReadySet tool in PHENIX. Model validation was performed using MolProbity (*77*) and EMRinger (*78*) in PHENIX, and relevant metrics are listed in table S1-3.

### Statistical analysis

Statistical analysis and curve fitting was carried out with the GraphPad Prism 10 software, and statistical significance was assigned to differences with a p value of <0.05. GraphPad Prism Michaelis-Menten non-linear regression analysis was used to fit data displayed in Figure 1B. This non-linear regression model is given by:

Y = V_max_*X/(K_m_+X), where V_max_ is the maximum velocity in the same unit as Y and K_m_ is the Michaelis-Menten constant, in the same units as X. K_m_ is the substrate concentration needed to achieve a half-maximum enzyme velocity.

ns: p ≥ 0.05. The number of replicates (n) used for calculating statistics is specified in the figure legends.

## Acknowledgments

We wish to thank Sara Abad Herrera, Bo Højen Justesen (Ruhr University Bochum) for helpful discussions. We thank Thomas Boesen, Andreas Bøggild, and Taner Drace for technical support during EM data collection at the EMBION Danish National cryo-EM facility of Aarhus University (5072-00025B, Danish Agency for Research and Higher Education) and Jesper Lykkegaard Karlsen for scientific computing support. We are also grateful to Malika Ould Ali and Laura Pieri for support during proteoliposome imaging at the I2BC cryo-EM platform. This work also benefited from the Cell and Tissue Imaging core facility (PICT IBiSA) at Institut Curie.

## Funding

Agence Nationale de la Recherche ANR-21-CE11-0015-01 (GL)

Lundbeck Foundation LF, R335-2019-2053 (JAL)

Deutsche Forschungsgemeinschaft GU 1133/13-1 (TGP)

Deutsche Forschungsgemeinschaft GU 1133/15-1 (TGP)

Independent Research Fund Denmark | Nature and Universe FNU, 1026-00024B (RLLM)

French National Research Infrastructure France-BioImaging ANR-10-INSB-04 (DL)

Region Ile de France Sesame 2018 3D EM/CLEM EXO039200 (DL)

French Infrastructure for Integrated Structural Biology (FRISBI) (ANR-10-INSB-05-05)

## Author contributions

Conceptualization: JAL, GL

Methodology: ABP, PF, CM, ADD, TD, DL, TGP, RLLM, JAL, GL

Validation: ABP, PF, DL, RLLM, JAL, GL

Formal analysis: ABP, PF, RLLM, JAL, GL

Investigation: ABP, PF, ADD, TD, RLLM

Resources: MMF, YI

Writing - Original Draft: ABP, PF, JAL, GL

Writing - Review & Editing: ABP, PF, TD, DL, TGP, RLLM, JAL, GL

Visualization: ABP, PF, RLLM, JAL, GL

Supervision: JAL, GL

Project administration: JAL, GL

Funding acquisition: TGP, DL, RLLM, JAL, GL

## Data and materials availability

All data are available in the main text or the supplementary materials.

## References

1. R. Panatala, H. Hennrich, J. C. M. Holthuis, Inner workings and biological impact of phospholipid flippases. J. Cell Sci. 128, 2021–2032 (2015).

2. J. H. Lorent, K. R. Levental, L. Ganesan, G. Rivera-Longsworth, E. Sezgin, M. Doktorova, E. Lyman, I. Levental, Plasma membranes are asymmetric in lipid unsaturation, packing and protein shape. Nat. Chem. Biol. 16, 644–652 (2020).

3. M. Murate, M. Abe, K. Kasahara, K. Iwabuchi, M. Umeda, T. Kobayashi, Transbilayer distribution of lipids at nano scale. J. Cell Sci. 128, 1627–1638 (2015).

4. J. Cerbón, V. Calderón, Changes of the compositional asymmetry of phospholipids associated to the increment in the membrane surface potential. Biochim. Biophys. Acta 1067, 139–144 (1991).

5. T. Tsuji, J. Cheng, T. Tatematsu, A. Ebata, H. Kamikawa, A. Fujita, S. Gyobu, K. Segawa, H. Arai, T. Taguchi, S. Nagata, T. Fujimoto, Predominant localization of phosphatidylserine at the cytoplasmic leaflet of the ER, and its TMEM16K-dependent redistribution. Proc. Natl. Acad. Sci. U. S. A. 116, 13368–13373 (2019).

6. J. C. M. Holthuis, A. K. Menon, Lipid landscapes and pipelines in membrane homeostasis. Nature 510, 48–57 (2014).

7. J. Bigay, B. Antonny, Curvature, lipid packing, and electrostatics of membrane organelles: defining cellular territories in determining specificity. Dev. Cell 23, 886–895 (2012).

8. R. L. López-Marqués, L. R. Poulsen, A. Bailly, M. Geisler, T. G. Pomorski, M. G. Palmgren, Structure and mechanism of ATP-dependent phospholipid transporters. Biochim. Biophys. Acta 1850, 461–475 (2015).

9. S. Bryde, H. Hennrich, P. M. Verhulst, P. F. Devaux, G. Lenoir, J. C. M. Holthuis, CDC50 proteins are critical components of the human class-1 P4-ATPase transport machinery. J. Biol. Chem. 285, 40562–40572 (2010).

10. G. Lenoir, P. Williamson, C. F. Puts, J. C. M. Holthuis, Cdc50p plays a vital role in the ATPase reaction cycle of the putative aminophospholipid transporter Drs2p. J. Biol. Chem. 284, 17956–17967 (2009).

11. K. Segawa, S. Kurata, S. Nagata, The CDC50A extracellular domain is required for forming a functional complex with and chaperoning phospholipid flippases to the plasma membrane. J. Biol. Chem. 293, 2172–2182 (2018).

12. M. Timcenko, J. A. Lyons, D. Januliene, J. J. Ulstrup, T. Dieudonné, C. Montigny, M.-R. Ash, J. L. Karlsen, T. Boesen, W. Kühlbrandt, G. Lenoir, A. Moeller, P. Nissen, Structure and autoregulation of a P4-ATPase lipid flippase. Nature 571, 366–370 (2019).

13. M. Hiraizumi, K. Yamashita, T. Nishizawa, O. Nureki, Cryo-EM structures capture the transport cycle of the P4-ATPase flippase. Science 365, 1149–1155 (2019).

14. L. Bai, A. Kovach, Q. You, H.-C. Hsu, G. Zhao, H. Li, Autoinhibition and activation mechanisms of the eukaryotic lipid flippase Drs2p-Cdc50p. Nat. Commun. 10, 4142 (2019).

15. H. Nakanishi, T. Nishizawa, K. Segawa, O. Nureki, Y. Fujiyoshi, S. Nagata, K. Abe, Transport Cycle of Plasma Membrane Flippase ATP11C by Cryo-EM. Cell Rep. 32, 108208 (2020).

16. J. A. Lyons, M. Timcenko, T. Dieudonné, G. Lenoir, P. Nissen, P4-ATPases: how an old dog learnt new tricks - structure and mechanism of lipid flippases. Curr. Opin. Struct. Biol. 63, 65–73 (2020).

17. O. E. Onat, S. Gulsuner, K. Bilguvar, A. Nazli Basak, H. Topaloglu, M. Tan, U. Tan, M. Gunel, T. Ozcelik, Missense mutation in the ATPase, aminophospholipid transporter protein ATP8A2 is associated with cerebellar atrophy and quadrupedal locomotion. Eur. J. Hum. Genet. EJHG 21, 281–285 (2013).

18. L. N. Bull, M. J. van Eijk, L. Pawlikowska, J. A. DeYoung, J. A. Juijn, M. Liao, L. W. Klomp, N. Lomri, R. Berger, B. F. Scharschmidt, A. S. Knisely, R. H. Houwen, N. B. Freimer, A gene encoding a P-type ATPase mutated in two forms of hereditary cholestasis. Nat. Genet. 18, 219–224 (1998).

19. L. Gao, M. J. Emond, T. Louie, C. Cheadle, A. E. Berger, N. Rafaels, C. Vergara, Y. Kim, M. A. Taub, I. Ruczinski, S. C. Mathai, S. S. Rich, D. A. Nickerson, L. K. Hummers, M. J. Bamshad, P. M. Hassoun, R. A. Mathias, National Heart, Lung, and Blood Institute GO Exome Sequencing Project, K. C. Barnes, Identification of Rare Variants in ATP8B4 as a Risk Factor for Systemic Sclerosis by Whole-Exome Sequencing. Arthritis Rheumatol. Hoboken NJ 68, 191–200 (2016).

20. H. Holstege, M. Hulsman, C. Charbonnier, B. Grenier-Boley, O. Quenez, D. Grozeva, J. G. J. van Rooij, R. Sims, S. Ahmad, N. Amin, P. J. Norsworthy, O. Dols-Icardo, H. Hummerich, A. Kawalia, P. Amouyel, G. W. Beecham, C. Berr, J. C. Bis, A. Boland, P. Bossù, F. Bouwman, J. Bras, D. Campion, J. N. Cochran, A. Daniele, J.-F. Dartigues, S. Debette, J.-F. Deleuze, N. Denning, A. L. DeStefano, L. A. Farrer, M. V. Fernández, N. C. Fox, D. Galimberti, E. Genin, J. J. P. Gille, Y. Le Guen, R. Guerreiro, J. L. Haines, C. Holmes, M. A. Ikram, M. K. Ikram, I. E. Jansen, R. Kraaij, M. Lathrop, A. W. Lemstra, A. Lleó, L. Luckcuck, M. M. A. M. Mannens, R. Marshall, E. R. Martin, C. Masullo, R. Mayeux, P. Mecocci, A. Meggy, M. O. Mol, K. Morgan, R. M. Myers, B. Nacmias, A. C. Naj, V. Napolioni, F. Pasquier, P. Pastor, M. A. Pericak-Vance, R. Raybould, R. Redon, M. J. T. Reinders, A.-C. Richard, S. G. Riedel-Heller, F. Rivadeneira, S. Rousseau, N. S. Ryan, S. Saad, P. Sanchez-Juan, G. D. Schellenberg, P. Scheltens, J. M. Schott, D. Seripa, S. Seshadri, D. Sie, E. A. Sistermans, S. Sorbi, R. van Spaendonk, G. Spalletta, N. Tesi, B. Tijms, A. G. Uitterlinden, S. J. van der Lee, P. J. Visser, M. Wagner, D. Wallon, L.-S. Wang, A. Zarea, J. Clarimon, J. C. van Swieten, M. D. Greicius, J. S. Yokoyama, C. Cruchaga, J. Hardy, A. Ramirez, S. Mead, W. M. van der Flier, C. M. van Duijn, J. Williams, G. Nicolas, C. Bellenguez, J.-C. Lambert, Exome sequencing identifies rare damaging variants in ATP8B4 and ABCA1 as risk factors for Alzheimer’s disease. Nat. Genet. 54, 1786–1794 (2022).

21. H.-W. Shin, H. Takatsu, Substrates, regulation, cellular functions, and disease associations of P4-ATPases. *Commun*. Biol. 8, 135 (2025).

22. H.-W. Shin, H. Takatsu, Substrates of P4-ATPases: beyond aminophospholipids (phosphatidylserine and phosphatidylethanolamine). FASEB J. Off. Publ. Fed. Am. Soc. Exp. Biol. 33, 3087–3096 (2019).

23. J. A. Coleman, M. C. M. Kwok, R. S. Molday, Localization, purification, and functional reconstitution of the P4-ATPase Atp8a2, a phosphatidylserine flippase in photoreceptor disc membranes. J. Biol. Chem. 284, 32670–32679 (2009).

24. J. Wang, L. L. Molday, T. Hii, J. A. Coleman, T. Wen, J. P. Andersen, R. S. Molday, Proteomic Analysis and Functional Characterization of P4-ATPase Phospholipid Flippases from Murine Tissues. Sci. Rep. 8, 10795 (2018).

25. L. R. Poulsen, R. L. López-Marqués, P. R. Pedas, S. C. McDowell, E. Brown, R. Kunze, J. F. Harper, T. G. Pomorski, M. Palmgren, A phospholipid uptake system in the model plant Arabidopsis thaliana. Nat. Commun. 6, 7649 (2015).

26. Y. Muranaka, R. Shigetomi, Y. Iwasaki, A. Hamamoto, K. Nakayama, H. Takatsu, H.-W. Shin, Novel phosphatidylinositol flippases contribute to phosphoinositide homeostasis in the plasma membrane. Biochem. J. 481, 1187–1202 (2024).

27. T. Dieudonné, S. A. Herrera, M. J. Laursen, M. Lejeune, C. Stock, K. Slimani, C. Jaxel, J. A. Lyons, C. Montigny, T. G. Pomorski, P. Nissen, G. Lenoir, Autoinhibition and regulation by phosphoinositides of ATP8B1, a human lipid flippase associated with intrahepatic cholestatic disorders. eLife 11, e75272 (2022).

28. T. Dieudonné, F. Kümmerer, M. J. Laursen, C. Stock, R. K. Flygaard, S. Khalid, G. Lenoir, J. A. Lyons, K. Lindorff-Larsen, P. Nissen, Activation and substrate specificity of the human P4-ATPase ATP8B1. Nat. Commun. 14, 7492 (2023).

29. X. Tang, M. S. Halleck, R. A. Schlegel, P. Williamson, A subfamily of P-type ATPases with aminophospholipid transporting activity. Science 272, 1495–1497 (1996).

30. M. Timcenko, T. Dieudonné, C. Montigny, T. Boesen, J. A. Lyons, G. Lenoir, P. Nissen, Structural Basis of Substrate-Independent Phosphorylation in a P4-ATPase Lipid Flippase. J. Mol. Biol., 167062 (2021).

31. H. Azouaoui, C. Montigny, T. Dieudonné, P. Champeil, A. Jacquot, J. L. Vázquez-Ibar, P. Le Maréchal, J. Ulstrup, M.-R. Ash, J. A. Lyons, P. Nissen, G. Lenoir, High phosphatidylinositol 4-phosphate (PI4P)-dependent ATPase activity for the Drs2p-Cdc50p flippase after removal of its N- and C-terminal extensions. J. Biol. Chem. 292, 7954–7970 (2017).

32. P. Natarajan, K. Liu, D. V. Patil, V. A. Sciorra, C. L. Jackson, T. R. Graham, Regulation of a Golgi flippase by phosphoinositides and an ArfGEF. Nat. Cell Biol. 11, 1421–1426 (2009).

33. H. Azouaoui, C. Montigny, M.-R. Ash, F. Fijalkowski, A. Jacquot, C. Grønberg, R. L. López-Marqués, M. G. Palmgren, M. Garrigos, M. le Maire, P. Decottignies, P. Gourdon, P. Nissen, P. Champeil, G. Lenoir, A high-yield co-expression system for the purification of an intact Drs2p-Cdc50p lipid flippase complex, critically dependent on and stabilized by phosphatidylinositol-4-phosphate. PloS One 9, e112176 (2014).

34. P. Natarajan, J. Wang, Z. Hua, T. R. Graham, Drs2p-coupled aminophospholipid translocase activity in yeast Golgi membranes and relationship to in vivo function. Proc. Natl. Acad. Sci. U. S. A. 101, 10614–10619 (2004).

35. X. Zhou, T. R. Graham, Reconstitution of phospholipid translocase activity with purified Drs2p, a type-IV P-type ATPase from budding yeast. Proc. Natl. Acad. Sci. U. S. A. 106, 16586–16591 (2009).

36. S. A. Herrera, B. H. Justesen, T. Dieudonné, C. Montigny, P. Nissen, G. Lenoir, T. Günther Pomorski, Direct evidence of lipid transport by the Drs2-Cdc50 flippase upon truncation of its terminal regions. Protein Sci. Publ. Protein Soc. 33, e4855 (2023).

37. K. Liu, K. Surendhran, S. F. Nothwehr, T. R. Graham, P4-ATPase requirement for AP-1/clathrin function in protein transport from the trans-Golgi network and early endosomes. Mol. Biol. Cell 19, 3526–3535 (2008).

38. W. E. Gall, N. C. Geething, Z. Hua, M. F. Ingram, K. Liu, S. I. Chen, T. R. Graham, Drs2p-dependent formation of exocytic clathrin-coated vesicles in vivo. Curr. Biol. CB 12, 1623– 1627 (2002).

39. L. Theorin, K. Faxén, D. M. Sørensen, R. Migotti, G. Dittmar, J. Schiller, D. L. Daleke, M. Palmgren, R. L. López-Marqués, T. Günther Pomorski, The lipid head group is the key element for substrate recognition by the P4 ATPase ALA2: a phosphatidylserine flippase. Biochem. J. 476, 783–794 (2019).

40. J. K. Paterson, K. Renkema, L. Burden, M. S. Halleck, R. A. Schlegel, P. Williamson, D. L. Daleke, Lipid specific activation of the murine P4-ATPase Atp8a1 (ATPase II). Biochemistry 45, 5367–5376 (2006).

41. A. K. Mandal, W. D. Cheung, J. M. Argüello, Characterization of a thermophilic P-type Ag+/Cu+-ATPase from the extremophile Archaeoglobus fulgidus. J. Biol. Chem. 277, 7201– 7208 (2002).

42. N. Alder-Baerens, Q. Lisman, L. Luong, T. Pomorski, J. C. M. Holthuis, Loss of P4 ATPases Drs2p and Dnf3p disrupts aminophospholipid transport and asymmetry in yeast post-Golgi secretory vesicles. Mol. Biol. Cell 17, 1632–1642 (2006).

43. M. Takar, Y. Wu, T. R. Graham, The Essential Neo1 Protein from Budding Yeast Plays a Role in Establishing Aminophospholipid Asymmetry of the Plasma Membrane. J. Biol. Chem. 291, 15727–15739 (2016).

44. M. S. Jensen, S. Costa, T. Günther-Pomorski, R. L. López-Marqués, Cell-Based Lipid Flippase Assay Employing Fluorescent Lipid Derivatives. Methods Mol. Biol. Clifton NJ 1377, 371–382 (2016).

45. J. A. Davis, R. B. Pares, T. Bernstein, S. C. McDowell, E. Brown, J. Stubrich, A. Rosenberg, E. B. Cahoon, R. E. Cahoon, L. R. Poulsen, M. Palmgren, R. L. López-Marqués, J. F. Harper, The Lipid Flippases ALA4 and ALA5 Play Critical Roles in Cell Expansion and Plant Growth. Plant Physiol. 182, 2111–2125 (2020).

46. R. L. López-Marqués, L. R. Poulsen, S. Hanisch, K. Meffert, M. J. Buch-Pedersen, M. K. Jakobsen, T. G. Pomorski, M. G. Palmgren, Intracellular targeting signals and lipid specificity determinants of the ALA/ALIS P4-ATPase complex reside in the catalytic ALA alpha-subunit. Mol. Biol. Cell 21, 791–801 (2010).

47. L. R. Poulsen, R. L. López-Marqués, S. C. McDowell, J. Okkeri, D. Licht, A. Schulz, T. Pomorski, J. F. Harper, M. G. Palmgren, The Arabidopsis P4-ATPase ALA3 localizes to the golgi and requires a beta-subunit to function in lipid translocation and secretory vesicle formation. Plant Cell 20, 658–676 (2008).

48. P. R. Cullis, M. J. Hope, M. B. Bally, T. D. Madden, L. D. Mayer, D. B. Fenske, Influence of pH gradients on the transbilayer transport of drugs, lipids, peptides and metal ions into large unilamellar vesicles. Biochim. Biophys. Acta 1331, 187–211 (1997).

49. B. Ploier, A. K. Menon, A Fluorescence-based Assay of Phospholipid Scramblase Activity. J. Vis. Exp. JoVE, 54635 (2016).

50. J. D. Brunner, S. Schenck, Preparation of Proteoliposomes with Purified TMEM16 Protein for Accurate Measures of Lipid Scramblase Activity. Methods Mol. Biol. Clifton NJ 1949, 181– 199 (2019).

51. M. Traïkia, D. E. Warschawski, O. Lambert, J.-L. Rigaud, P. F. Devaux, Asymmetrical membranes and surface tension. Biophys. J. 83, 1443–1454 (2002).

52. S. Shukla, T. Baumgart, Enzymatic trans-bilayer lipid transport: Mechanisms, efficiencies, slippage, and membrane curvature. Biochim. Biophys. Acta Biomembr. 1863, 183534 (2021).

53. A. L. Vestergaard, J. A. Coleman, T. Lemmin, S. A. Mikkelsen, L. L. Molday, B. Vilsen, R. S. Molday, M. Dal Peraro, J. P. Andersen, Critical roles of isoleucine-364 and adjacent residues in a hydrophobic gate control of phospholipid transport by the mammalian P4-ATPase ATP8A2. Proc. Natl. Acad. Sci. U. S. A. 111, E1334–1343 (2014).

54. X. Zhou, T. T. Sebastian, T. R. Graham, Auto-inhibition of Drs2p, a yeast phospholipid flippase, by its carboxyl-terminal tail. J. Biol. Chem. 288, 31807–31815 (2013).

55. F. Tadini-Buoninsegni, S. A. Mikkelsen, L. S. Mogensen, R. S. Molday, J. P. Andersen, Phosphatidylserine flipping by the P4-ATPase ATP8A2 is electrogenic. Proc. Natl. Acad. Sci. U. S. A. 116, 16332–16337 (2019).

56. D. Drew, R. A. North, K. Nagarathinam, M. Tanabe, Structures and General Transport Mechanisms by the Major Facilitator Superfamily (MFS). Chem. Rev. 121, 5289–5335 (2021).

57. J. A. Lyons, J. L. Parker, N. Solcan, A. Brinth, D. Li, S. T. A. Shah, M. Caffrey, S. Newstead, Structural basis for polyspecificity in the POT family of proton-coupled oligopeptide transporters. EMBO Rep. 15, 886–893 (2014).

58. W. M. Moore, R. J. Brea, C. Knittel, E. Wrightsman, B. Hui, J. Lou, C. F. Ancajas, M. D. Best, N. K. Devaraj, I. Budin, Leaflet specific phospholipid imaging using genetically encoded proximity sensors. BioRxiv Prepr. Serv. Biol., 2024.05.01.592120 (2025).

59. Y. Posor, W. Jang, V. Haucke, Phosphoinositides as membrane organizers. Nat. Rev. Mol. Cell Biol. 23, 797–816 (2022).

60. T. R. Graham, C. G. Burd, Coordination of Golgi functions by phosphatidylinositol 4-kinases. Trends Cell Biol. 21, 113–121 (2011).

61. G. R. V. Hammond, J. E. Burke, Novel roles of phosphoinositides in signaling, lipid transport, and disease. Curr. Opin. Cell Biol. 63, 57–67 (2020).

62. V. A. Sciorra, A. Audhya, A. B. Parsons, N. Segev, C. Boone, S. D. Emr, Synthetic genetic array analysis of the PtdIns 4-kinase Pik1p identifies components in a Golgi-specific Ypt31/rab-GTPase signaling pathway. Mol. Biol. Cell 16, 776–793 (2005).

63. C. F. Puts, G. Lenoir, J. Krijgsveld, P. Williamson, J. C. M. Holthuis, A P4-ATPase protein interaction network reveals a link between aminophospholipid transport and phosphoinositide metabolism. J. Proteome Res. 9, 833–842 (2010).

64. Z. Szentpetery, P. Várnai, T. Balla, Acute manipulation of Golgi phosphoinositides to assess their importance in cellular trafficking and signaling. Proc. Natl. Acad. Sci. U. S. A. 107, 8225–8230 (2010).

65. Y. J. Wang, J. Wang, H. Q. Sun, M. Martinez, Y. X. Sun, E. Macia, T. Kirchhausen, J. P. Albanesi, M. G. Roth, H. L. Yin, Phosphatidylinositol 4 phosphate regulates targeting of clathrin adaptor AP-1 complexes to the Golgi. Cell 114, 299–310 (2003).

66. C. Y. Chen, M. F. Ingram, P. H. Rosal, T. R. Graham, Role for Drs2p, a P-type ATPase and potential aminophospholipid translocase, in yeast late Golgi function. J. Cell Biol. 147, 1223–1236 (1999).

67. L. Wang, Y. Iwasaki, K. K. Andra, K. Pandey, A. K. Menon, P. Bütikofer, Scrambling of natural and fluorescently tagged phosphatidylinositol by reconstituted G protein-coupled receptor and TMEM16 scramblases. J. Biol. Chem. 293, 18318–18327 (2018).

68. P. M. Burgers, Overexpression of multisubunit replication factors in yeast. Methods San Diego Calif 18, 349–355 (1999).

69. Z. Hua, P. Fatheddin, T. R. Graham, An essential subfamily of Drs2p-related P-type ATPases is required for protein trafficking between Golgi complex and endosomal/vacuolar system. Mol. Biol. Cell 13, 3162–3177 (2002).

70. R. D. Gietz, R. A. Woods, Transformation of yeast by lithium acetate/single-stranded carrier DNA/polyethylene glycol method. Methods Enzymol. 350, 87–96 (2002).

71. A. Punjani, J. L. Rubinstein, D. J. Fleet, M. A. Brubaker, cryoSPARC: algorithms for rapid unsupervised cryo-EM structure determination. Nat. Methods 14, 290–296 (2017).

72. A. Punjani, H. Zhang, D. J. Fleet, Non-uniform refinement: adaptive regularization improves single-particle cryo-EM reconstruction. Nat. Methods 17, 1214–1221 (2020).

73. R. T. Kidmose, J. Juhl, P. Nissen, T. Boesen, J. L. Karlsen, B. P. Pedersen, Namdinator - automatic molecular dynamics flexible fitting of structural models into cryo-EM and crystallography experimental maps. IUCrJ 6, 526–531 (2019).

74. T. I. Croll, ISOLDE: a physically realistic environment for model building into low-resolution electron-density maps. Acta Crystallogr. Sect. Struct. Biol. 74, 519–530 (2018).

75. P. V. Afonine, B. P. Klaholz, N. W. Moriarty, B. K. Poon, O. V. Sobolev, T. C. Terwilliger, P. D. Adams, A. Urzhumtsev, New tools for the analysis and validation of cryo-EM maps and atomic models. Acta Crystallogr. Sect. Struct. Biol. 74, 814–840 (2018).

76. D. Liebschner, P. V. Afonine, M. L. Baker, G. Bunkóczi, V. B. Chen, T. I. Croll, B. Hintze, L. W. Hung, S. Jain, A. J. McCoy, N. W. Moriarty, R. D. Oeffner, B. K. Poon, M. G. Prisant, R. J. Read, J. S. Richardson, D. C. Richardson, M. D. Sammito, O. V. Sobolev, D. H. Stockwell, T. C. Terwilliger, A. G. Urzhumtsev, L. L. Videau, C. J. Williams, P. D. Adams, Macromolecular structure determination using X-rays, neutrons and electrons: recent developments in Phenix. Acta Crystallogr. Sect. Struct. Biol. 75, 861–877 (2019).

77. C. J. Williams, J. J. Headd, N. W. Moriarty, M. G. Prisant, L. L. Videau, L. N. Deis, V. Verma, D. A. Keedy, B. J. Hintze, V. B. Chen, S. Jain, S. M. Lewis, W. B. Arendall, J. Snoeyink, P. D. Adams, S. C. Lovell, J. S. Richardson, D. C. Richardson, MolProbity: More and better reference data for improved all-atom structure validation. Protein Sci. Publ. Protein Soc. 27, 293–315 (2018).

78. B. A. Barad, N. Echols, R. Y.-R. Wang, Y. Cheng, F. DiMaio, P. D. Adams, J. S. Fraser, EMRinger: side chain-directed model and map validation for 3D cryo-electron microscopy. Nat. Methods 12, 943–946 (2015).

