## Supplementary material for "Revisiting flippase specificity: Drs2-Cdc50 transports multiple anionic lipid substrates"

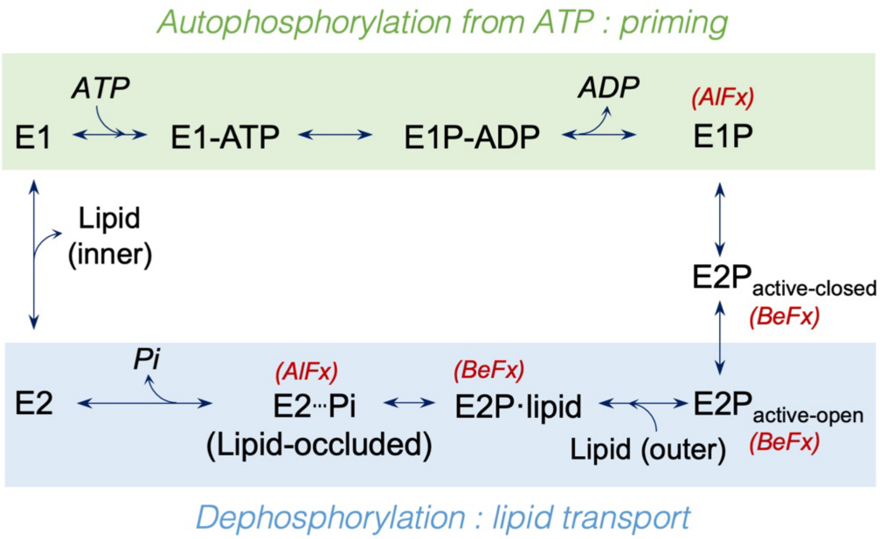

**Fig. S1: Post-Albers P4-ATPase catalytic cycle**

The Post-Albers P4-ATPase catalytic cycle includes the substrate-independent phosphorylation step from ATP (priming) and the substrate-dependent dephosphorylation (transport). The inhibitors used to capture the various catalytic states are indicated in red. The step at which the transported lipid is released toward the cytosolic leaflet has not yet been identified, and is highlighted by question mark.

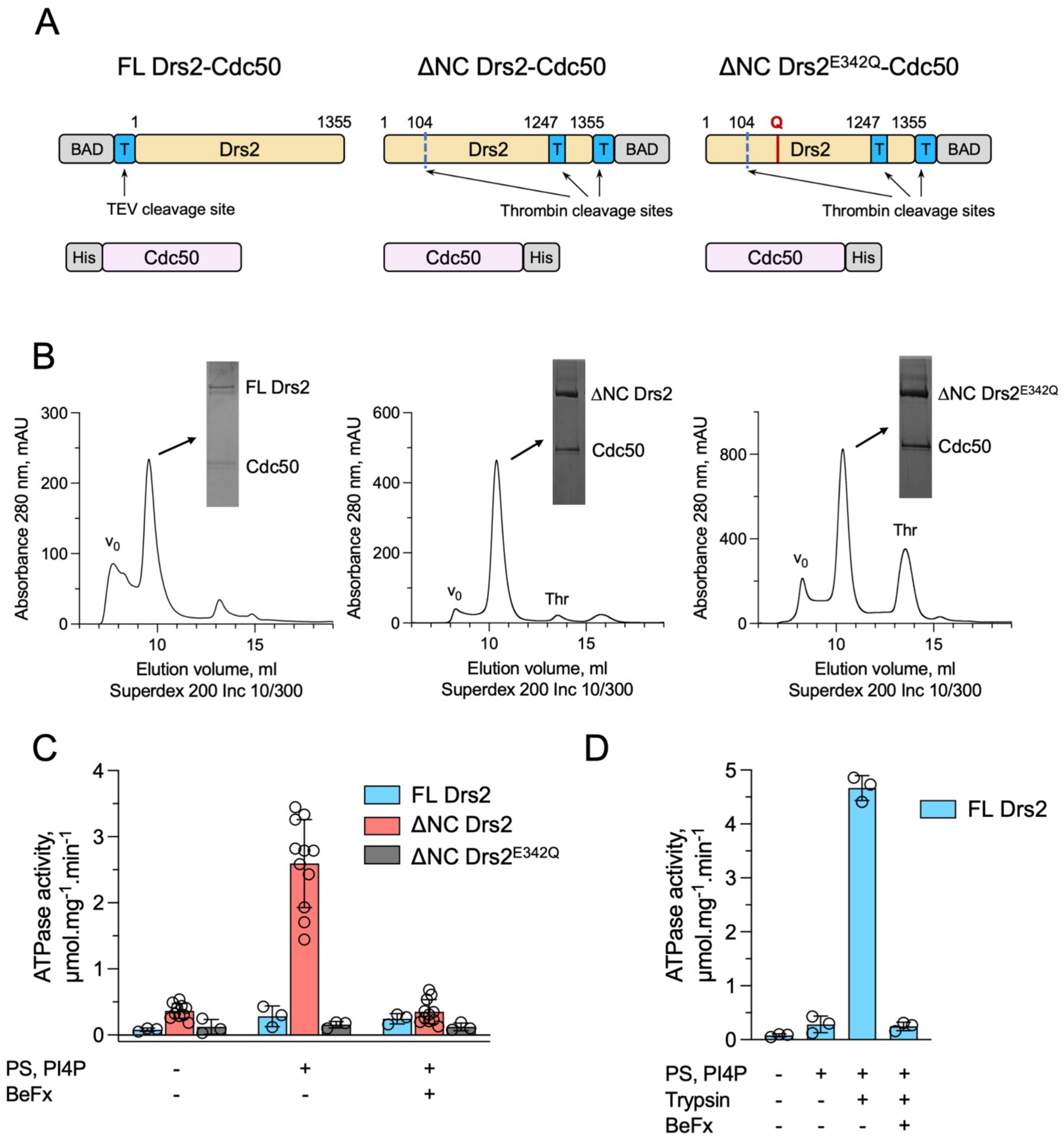

**Fig. S2: Protein purification and functional characterization**

**(A)**, Constructs used for co-overexpression of Drs2 and Cdc50 in yeast *S. cerevisiae*. Full-length (FL) Drs2 is tagged with a TEV-cleavable biotin acceptor domain (BAD) and Cdc50 is tagged with Histidine. The ΔNC Drs2 construct has a thrombin (Thr) cleavage site after residue 104 and an additional engineered thrombin cleavage site introduced in the C-terminus after residue 1247. The ΔNC E342Q Drs2 construct has a glutamate-to-glutamine substitution in the A-domain. **(B)**, Representative size exclusion chromatography (SEC) profiles of FL Drs2-Cdc50, ΔNC Drs2-

Cdc50 and ΔNC E342Q Drs2-Cdc50 complexes. Peak fractions were analyzed by Coomassie stained SDS-PAGE. $V_0$ – void volume, Thr – Thrombin. **(C)**, Specific ATPase activity of the indicated SEC-purified protein complexes in the absence and presence of PS, PI4P, and BeFx. **(D)**, ATPase activity of FL Drs2 without and with PS and PI4P, and after trypsin and BeFx addition. In panels C and D, ATPase activity was measured by an enzyme-coupled assay, as described in Methods. Bars show the mean ± SD from 3-11 measurements; individual data points shown.

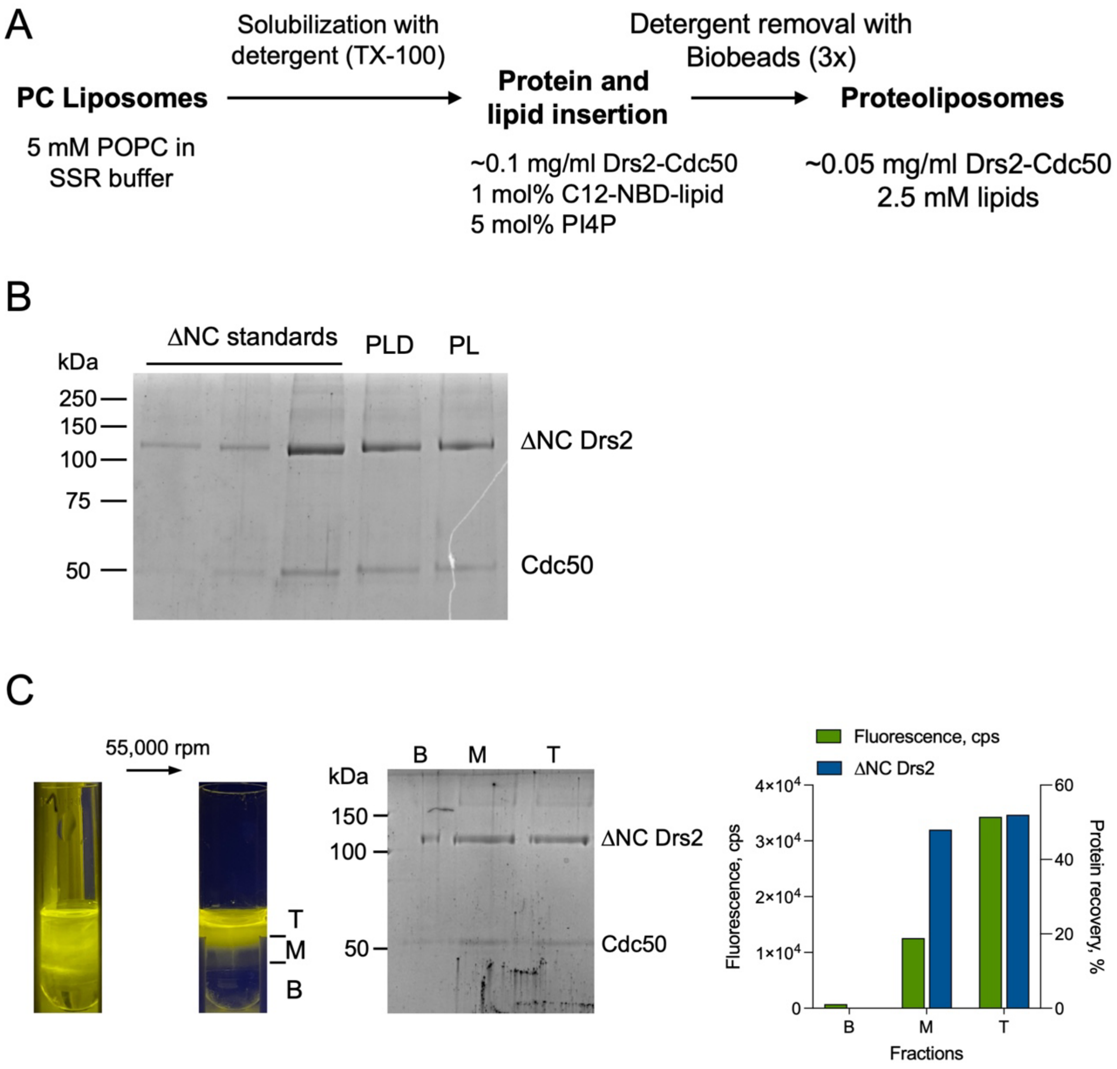

**Fig. S3: Reconstitution and functional analysis of ΔNC Drs2-Cdc50 in POPC liposomes**
**(A)**, Schematic overview of the reconstitution protocol used to incorporate Drs2-Cdc50 complexes into POPC liposomes. **(B)**, Protein recovery before (PLD) and after (PL) detergent removal, assessed by Coomassie blue-stained SDS-PAGE, showed minimal protein loss. **(C)**, Flotation assay of proteoliposomes containing PI4P and NBD-PS showed successful incorporation of ΔNC Drs2-Cdc50. Images (left) of the gradient before and after ultracentrifugation visually confirmed proteoliposome flotation and proper separation into bottom (B), middle (M), and top (T) fractions. Protein (Coomassie blue-stained SDS-PAGE, middle) and lipid (NBD fluorescence) were enriched in the middle (M) fraction and top (T) fraction.

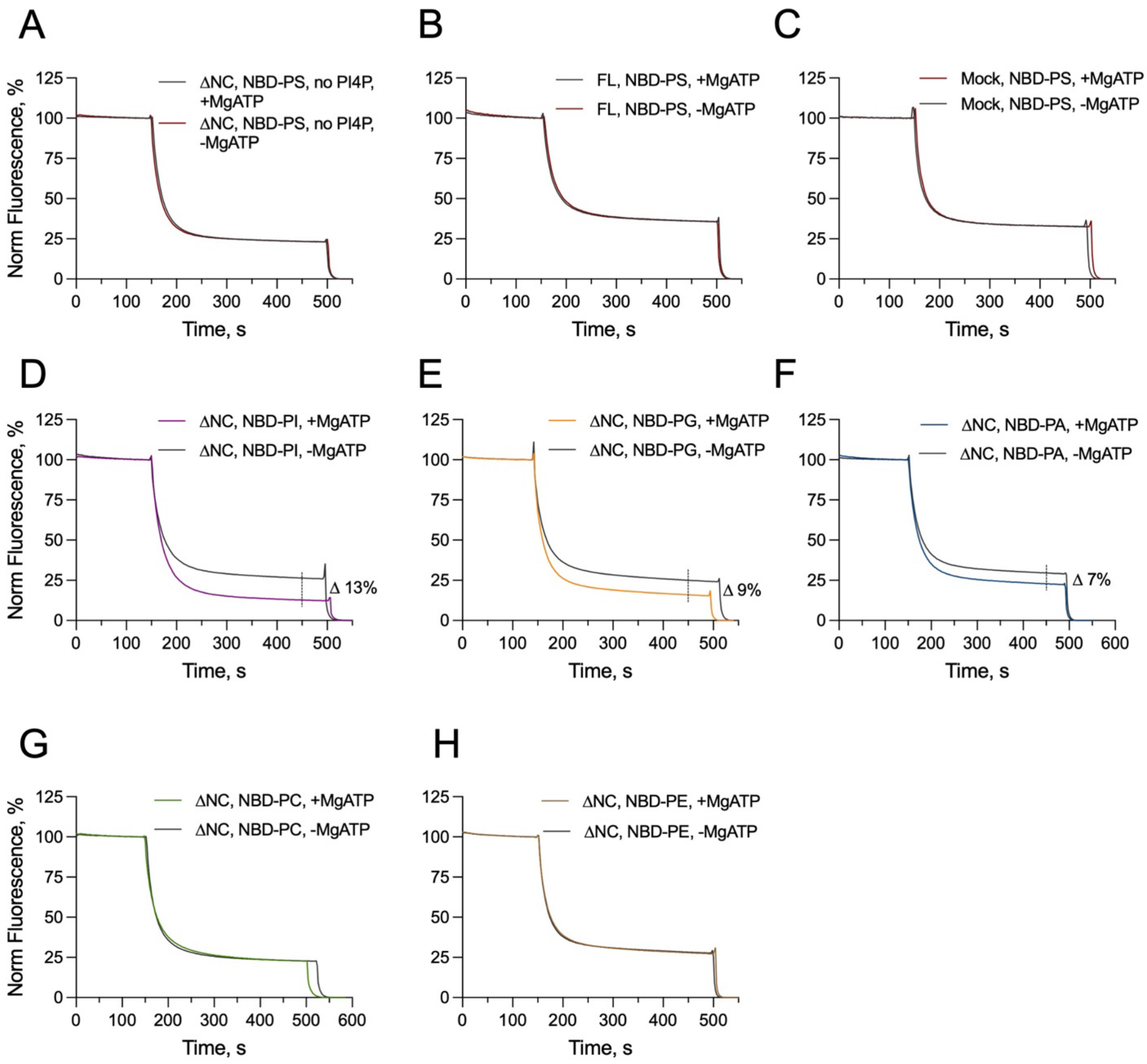

**Fig. S4: Drs2-Cdc50 mediated lipid transport in proteoliposomes**
Representative fluorescence traces of proteoliposomes reconstituted with 1 mol% NBD-lipid, 5 mol% PI4P and either ΔNC Drs2-Cdc50, Full-length (FL) Drs2-Cdc50, or without protein (mock). Samples were incubated for 30 min at 30 °C with or without 5 mM MgATP. Dithionite was added at ~150 s to quench outer leaflet fluorescence; TX-100 at ~500 s was added to disrupt all vesicles and quench the remaining inner leaflet NBD-lipids.

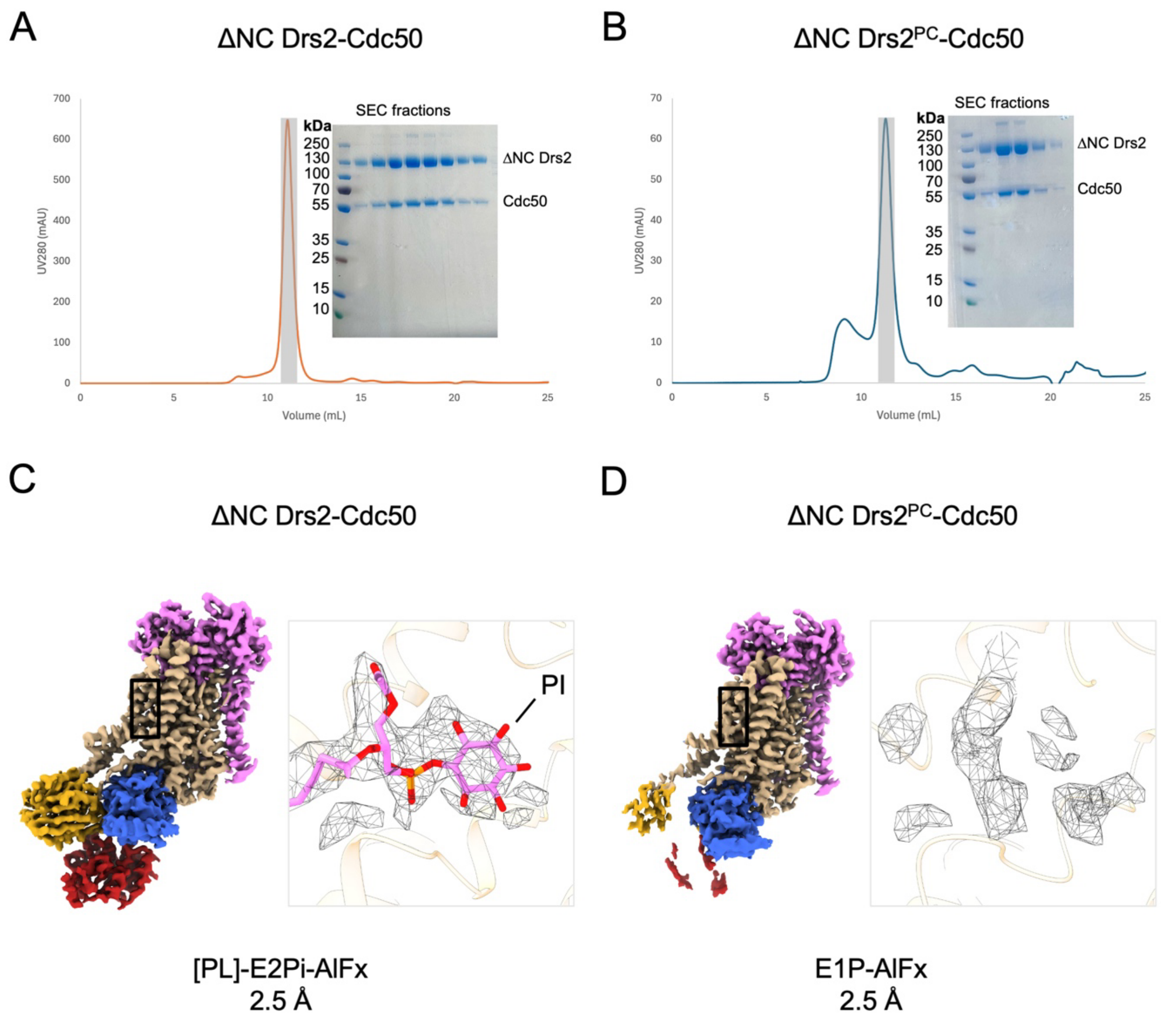

**Fig. S5: Purification and lipid exchange of ΔNC Drs2-Cdc50**

**(A)**, SEC profiles of ΔNC Drs2-Cdc50 and SDS-PAGE analysis of the protein peak. SEC was performed on a Superdex 300 increase 10/300 with an elution buffer of SSR containing 0.03 mg/mL LMNG and no glycerol. The ΔNC Drs2-Cdc50 complex eluted at a volume of 11.3 mL. The protein peak is shown as a grey shaded area. Bands corresponding to ΔNC Drs2 and Cdc50 are indicated. **(B)**, SEC profiles of ΔNC Drs2$^{PC}$-Cdc50 and SDS-PAGE analysis of the protein peak. SEC was performed on a Superdex 300 increase 10/300 with an elution buffer of SSR containing 0.03 mg/mL LMNG and no glycerol. The ΔNC Drs2$^{PC}$-Cdc50 complex eluted at a volume of 11.2 mL. The protein peak is shown as a grey shaded area. Bands corresponding to ΔNC Drs2 and Cdc50 are indicated. **(C)**, Cryo-EM map of ΔNC Drs2-Cdc50 in the E2Pi lipid occluded state, with a cryo-EM map density for a co-purified phospholipid consistent with PI (contour map/lipid density: 6.5/4.3). The model of [PI]-E2Pi-AlFx is shown here. **(D)**, Cryo-EM map of ΔNC Drs2$^{PC}$-Cdc50 in E1P conformation. No map density consistent with a phospholipid was present (contour map/lipid density: 8.3/6.5).

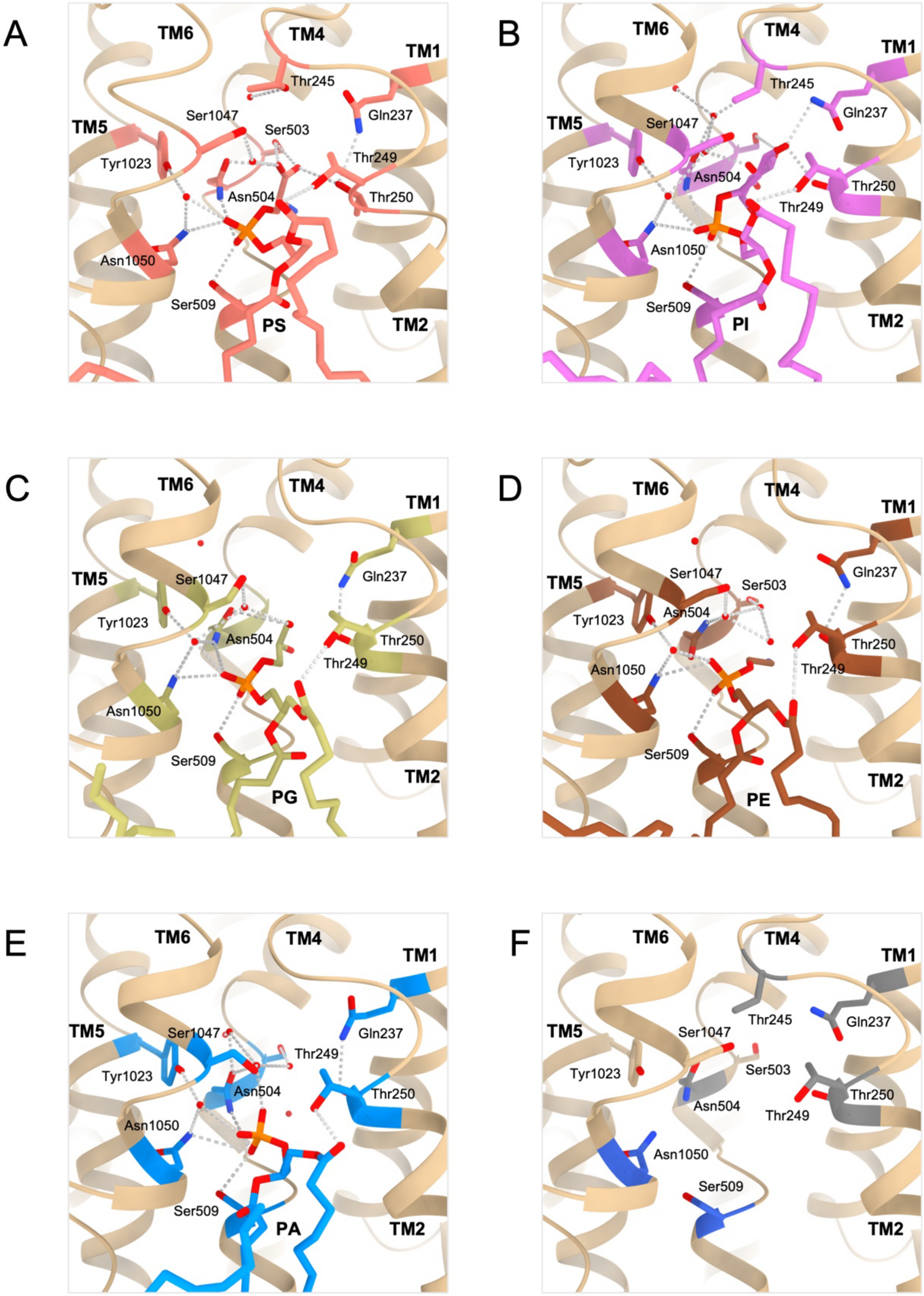

**Fig. S6: The hydrogen bonding network in the lipid occluded binding pocket**
**(A-E)**, The hydrogen bonding network to the substrate lipid. The lipids and respective interacting residues are colored as follow: PA = blue, PG = yellow, PI = pink, PE = brown and PS = salmon.

**(F)**, The conservation of the binding pocket highlighting the residues in direct and water-mediated contact with the substrate. Residues conserved across all human and yeast flippases are colored in blue. The residues conserved within the ATP8A family and Drs2 are colored in grey. Non-conserved residues involved in substrate contact are colored in tan.

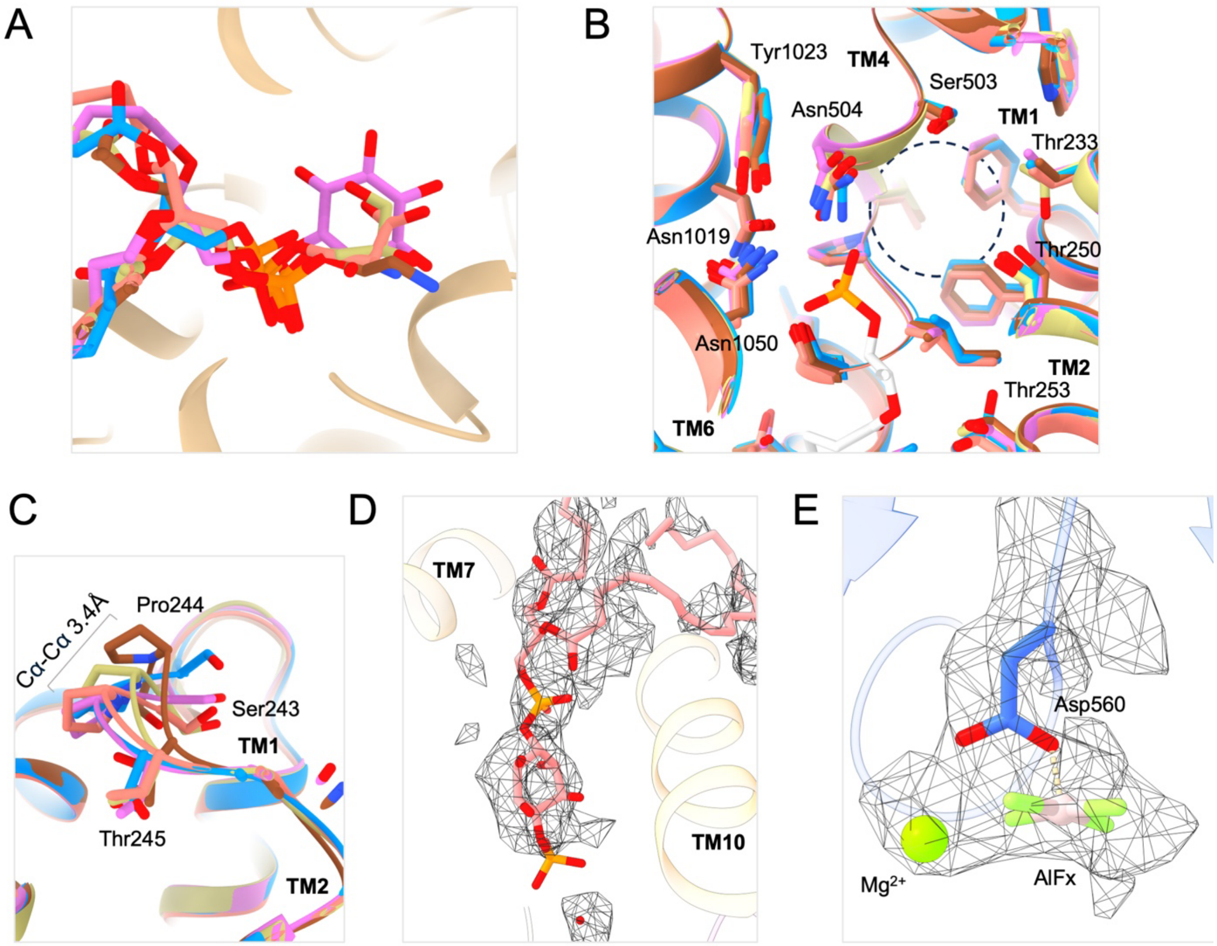

**Fig. S7: The lipid occluded model**

**(A)**, Comparison of the lipid binding mode of PA, PG, PI, PE and PS based on a superposition of TM7-10. PA = blue, PG = yellow, PI = pink, PE = brown and PS = salmon. Lipid binding is centered on the phosphate moiety, while the various headgroups share a common molecular backbone. **(B)**, Superposition of PA, PG, PI, PE and PS occluded states of Drs2 highlighting the largely invariable lipid binding pocket. The PI lipid without a headgroup is shown for reference. A dashed black cycle indicated the lipid headgroup binding cavity. Residues are colored according to substrate colors in panel A. **(C)**, Change in position of the loop region between TM1 and TM2. The models are colored according to substrate colors in panel A. The Cα-Cα distance between PA and PE is 3.4 Å. **(D)**, The PI4P binding site between TM7 and TM10. Model and cryo-EM map is for [PI]-E2Pi-AlFx. Contour map: 7.2. **(E)**, The phosphorylation site Asp560 coordinating AlFx and a $Mg^{2+}$ ion. Asp560 of the P-domain and AlFx are shown in sticks with the $Mg^{2+}$ ion as a green sphere. Model and cryo-EM map is for [PI]-E2Pi-AlFx. Contour map: 13.5.

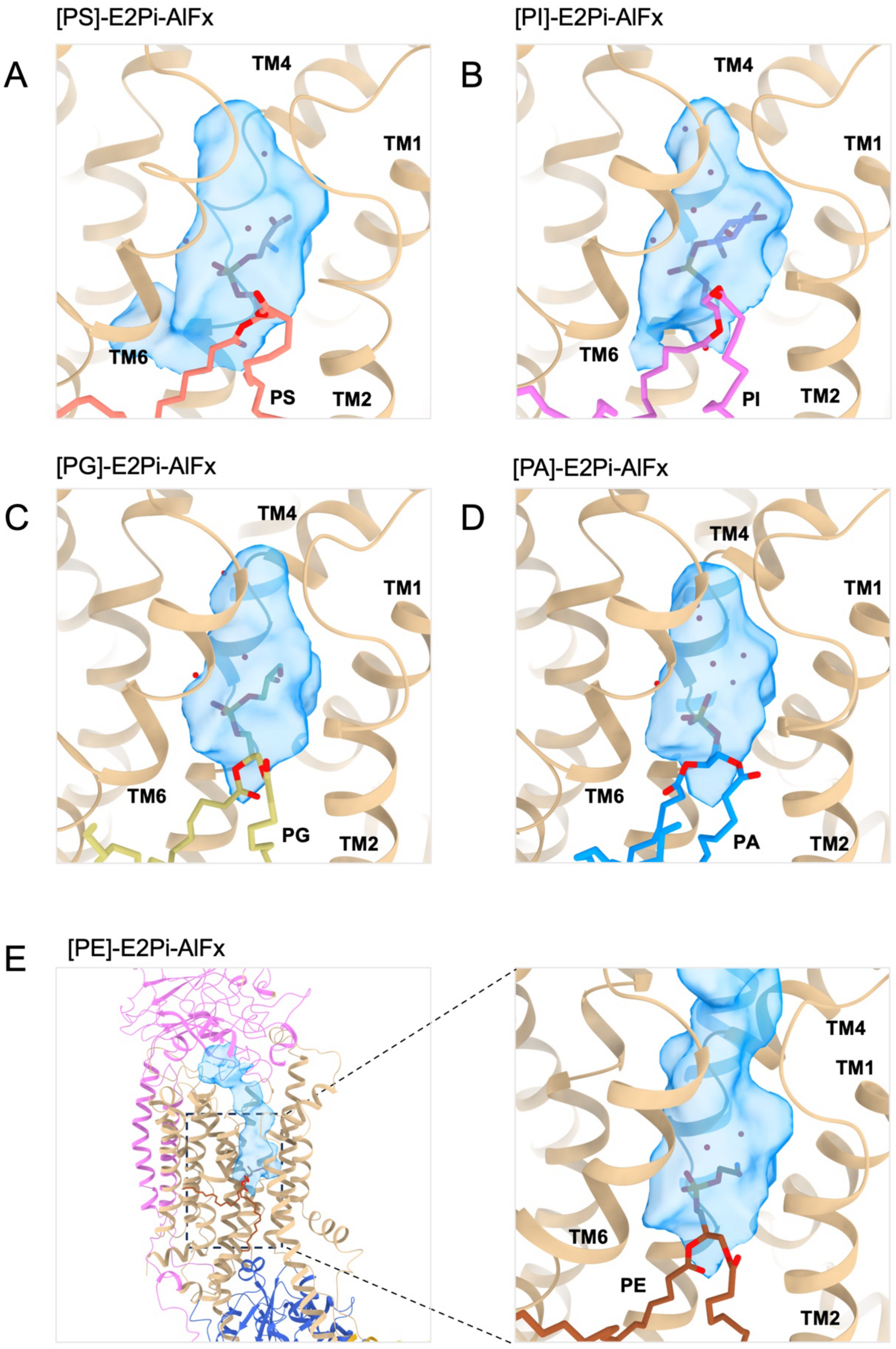

[PS]-E2Pi-AlFx
A
TM4
TM1
TM6
PS
TM2
[PI]-E2Pi-AlFx
B
TM4
TM1
TM6
PI
TM2
[PG]-E2Pi-AlFx
C
TM4
TM1
TM6
PG
TM2
[PA]-E2Pi-AlFx
D
TM4
TM1
TM6
PA
TM2
E
[PE]-E2Pi-AlFx
TM4
TM1
TM6
PE
TM2

**Fig. S8: Lipid headgroups are occluded in an invariant binding pocket**

The binding pocket is displayed for **(A)**, PS, **(B)**, PI, **(C)**, PG, **(D)**, PA and **(E)**, PE lipid occluded states. Cavities were calculated by the pyKVFinder tool in ChimeraX-1.9 with a probe radius of 1.4 Å and displayed as a transparent blue surface (*78*). The pocket maintains similar dimensions for PS, PI, PG and PA. For PE, the TM1-2 loop is partially occluding the substrate resulting in a continuous channel connecting to the luminal solvent.

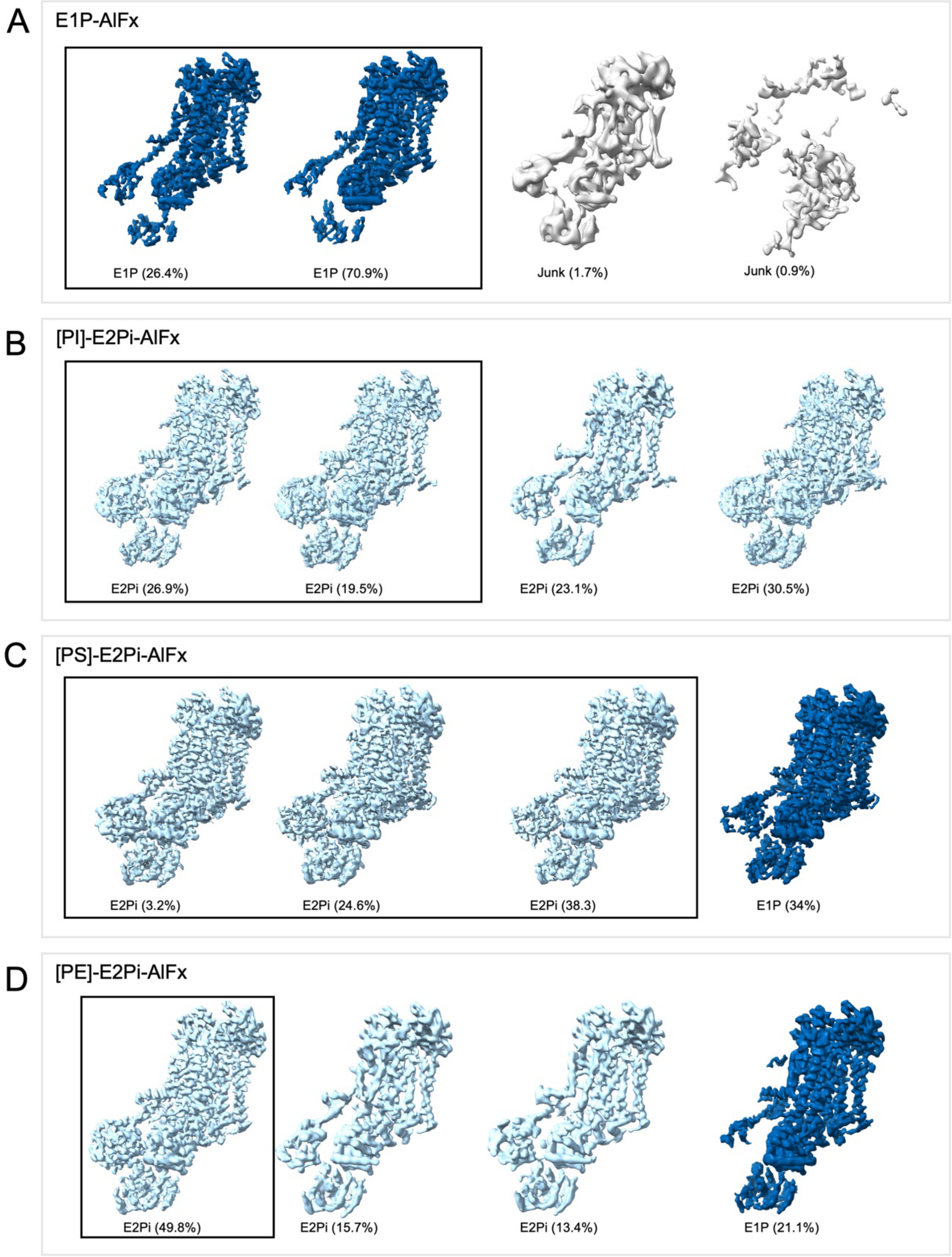

**Fig. S9: 3D classification reveals different conformations**

3D classification output from cryoSPARC of Drs2$^{PC}$-Cdc50 for various datasets: **(A)**, E1P-AlFx, **(B)**, [PI]-E2P-AlFx, **(C)**, [PS]-E2P-AlFx and **(D)**, [PE]-E2P-AlFx, indicating the relative conformational heterogeneity in the datasets. The distribution percentages of particles in each class are indicated. The volumes are colored by state – E1P = dark blue, E2Pi = light blue, junk = grey. The classes pooled for the final Non-Uniform Refinement are indicated by a black box.

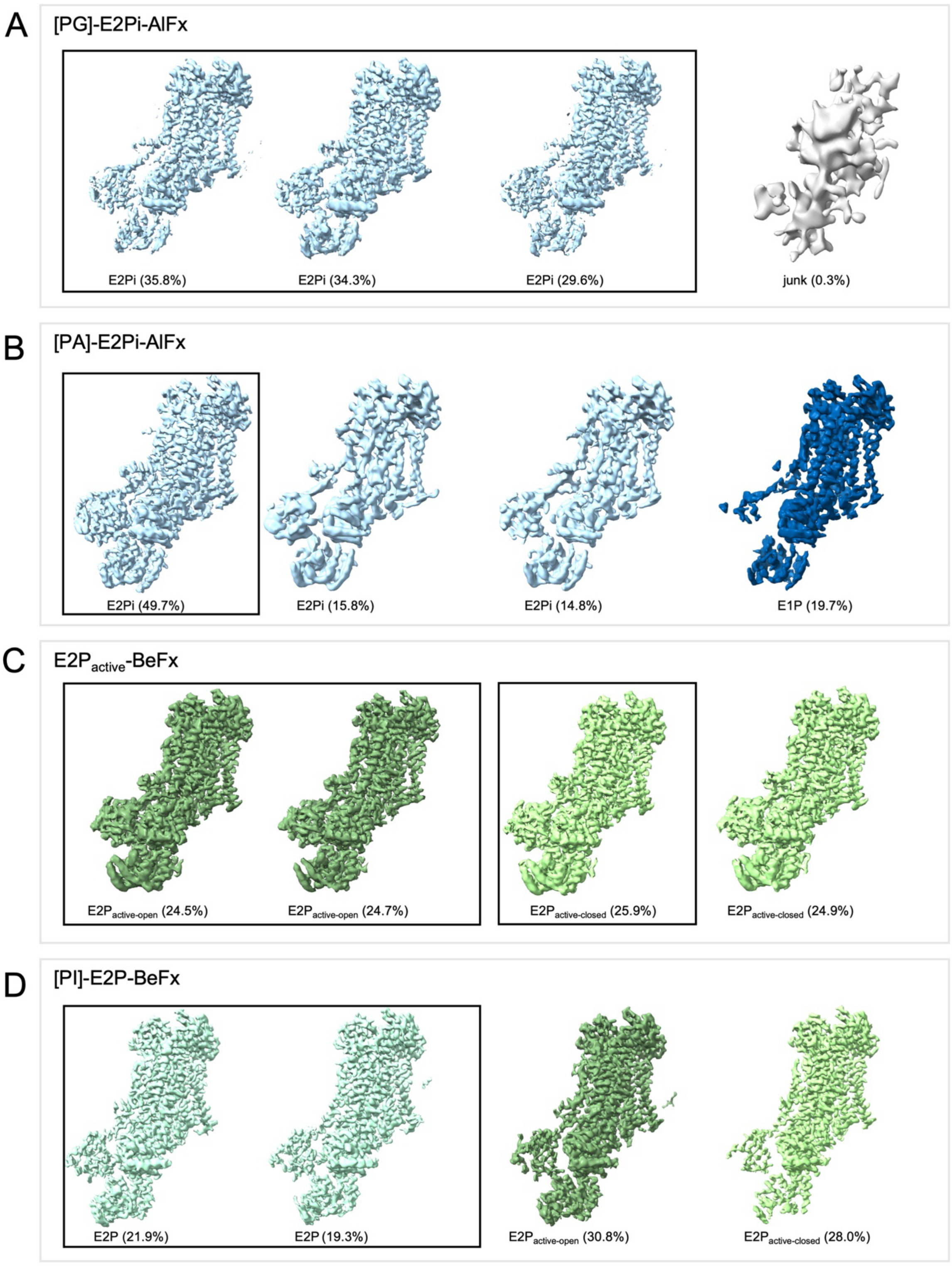

**Fig. S10: 3D classification reveals different conformations**

3D classification output from cryoSPARC of Drs2$^{PC}$-Cdc50 for various datasets: **(A)**, [PG]-E2P-AlFx, **(B)**, [PA]-E2P-AlFx, **(C)**, E2P$_{active}$-BeFx and **(D)**, [PI]-E2P-BeFx, indicating the relative conformational heterogeneity in the datasets. The distribution percentages of particles in each class are indicated. Volumes are colored by state – E1P = dark blue, E2Pi = light blue, dark green = E2P$_{active\text{-}open}$, light green = E2P$_{active\text{-}closed}$, E2P = aquamarine and junk = grey. The classes pooled for the final Non-Uniform Refinement are indicated by a black box.

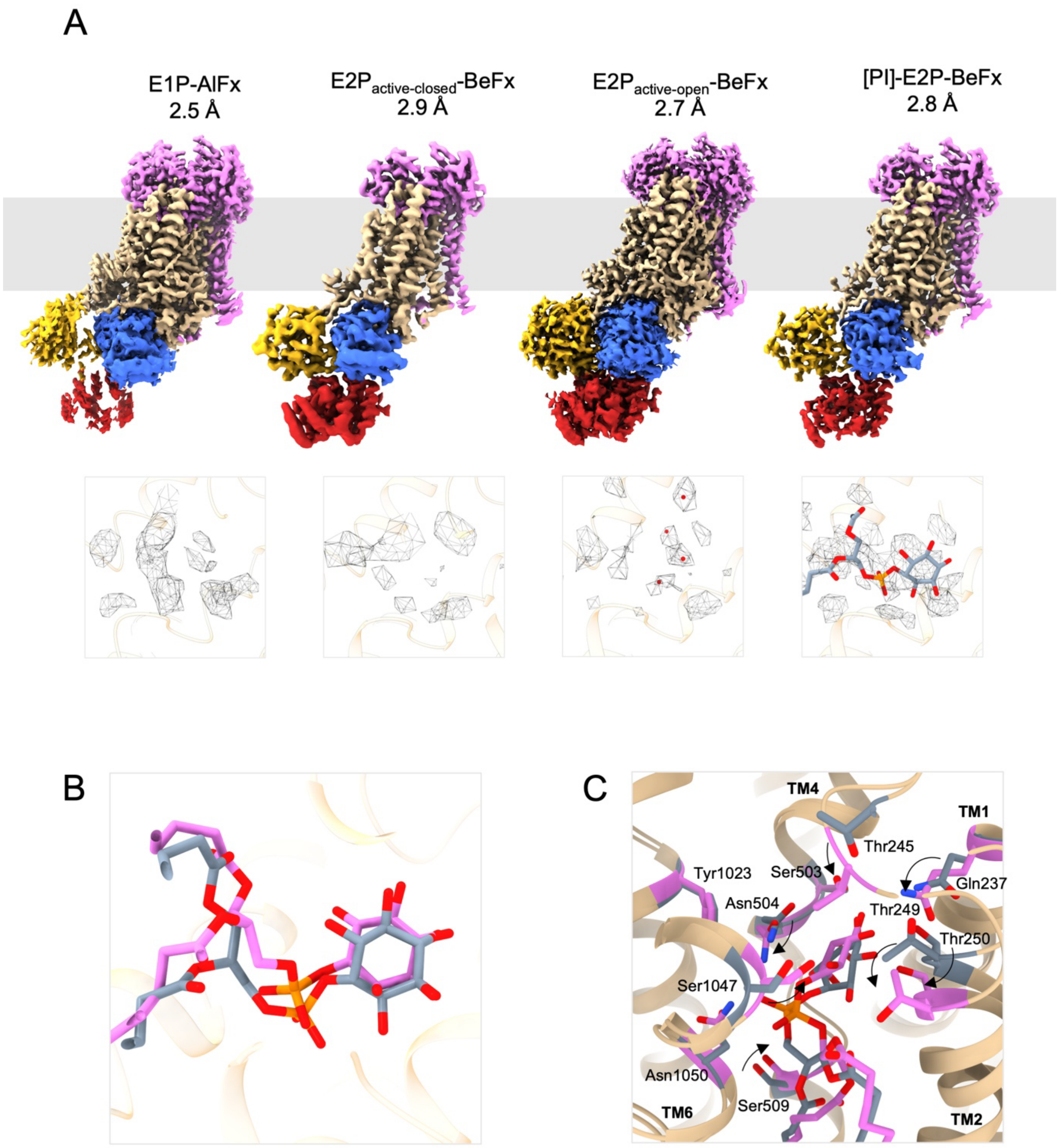

**Fig. S11: Cryo-EM maps of the Drs2-Cdc50 flippase complex inhibited by AlFx and BeFx** **(A)**, Cryo-EM maps of Drs2$^{\text{PC}}$-Cdc50 in the E1P and various E2P states. The maps are colored to highlight key subunits and domains; Cdc50 = pink, TMs = beige, P-domain = blue, A-domain = yellow and N-domain = red. The identity of the protein state, presence of occluded lipid and map resolution are indicated. Inset: Cryo-EM map density of the lipid binding pocket in E1P, E2P$_{\text{active-closed}}$, E2P$_{\text{active-open}}$ and [PI]-E2P-BeFx. Only [PI]-E2P-BeFx has recognizable cryo-EM map density for a lipid. The contour of the map vs contour used for visualization of the lipid and water densities: E1P (8.3 vs 6.5), E2P$_{\text{active-closed}}$ (10.8 vs 7.0), E2P$_{\text{active-open}}$ (5.89 vs 4.3), [PI]-E2P-BeFx

(8.0 vs 5.63). **(B)**, Comparison of the lipid binding mode of [PI]-E2P-BeFx (grey) and [PI]-E2Pi-AlFx (pink) based on a superposition of TM7-10. The lipid binding is centered on the phosphate moiety. **(C)**, Comparison of the residue movement during lipid binding (grey) and lipid occlusion (pink). Upon occlusion of the substrate, key residues are moving into hydrogen bonding distance of PI.

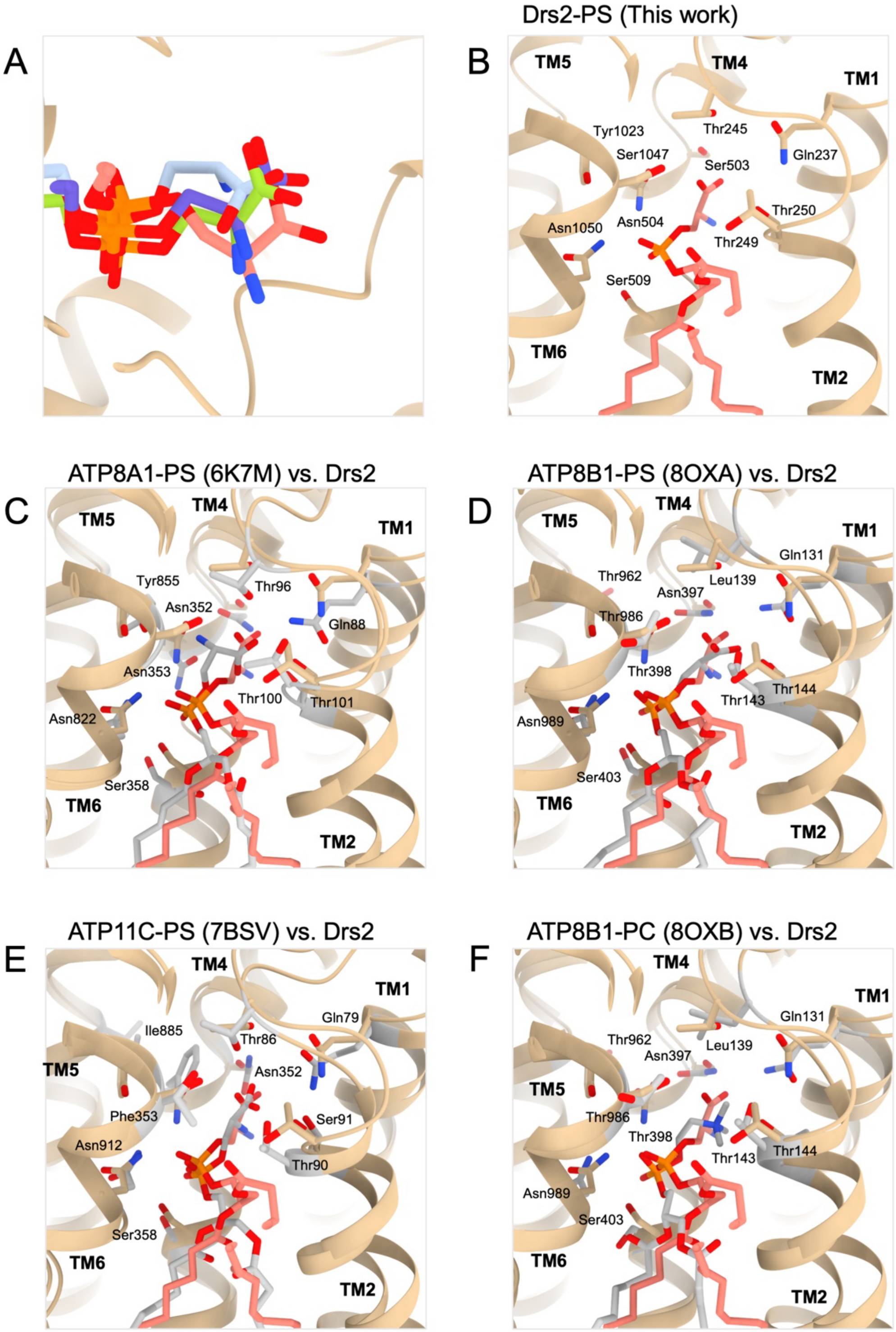

**Fig. S12: Comparison of PS-occluded models**

**(A)**, Comparison of the lipid binding mode of PS based on a superposition of TM7-10 in Drs2 (salmon), ATP8A1 (light blue), ATP8B1 (green) and ATP11C (purple). The PS binding is centered on the phosphate moiety, while the serine moiety varies. **(B)**, The lipid binding pocket of

Drs2 with the key interaction residues highlighted. **(C-F)**, The lipid binding pocket of ATP8A1-PS, ATP8B1-PS and ATP11C-PS, ATP8B1-PC respectively. The residues highlighted correspond to the ones shown in (B).

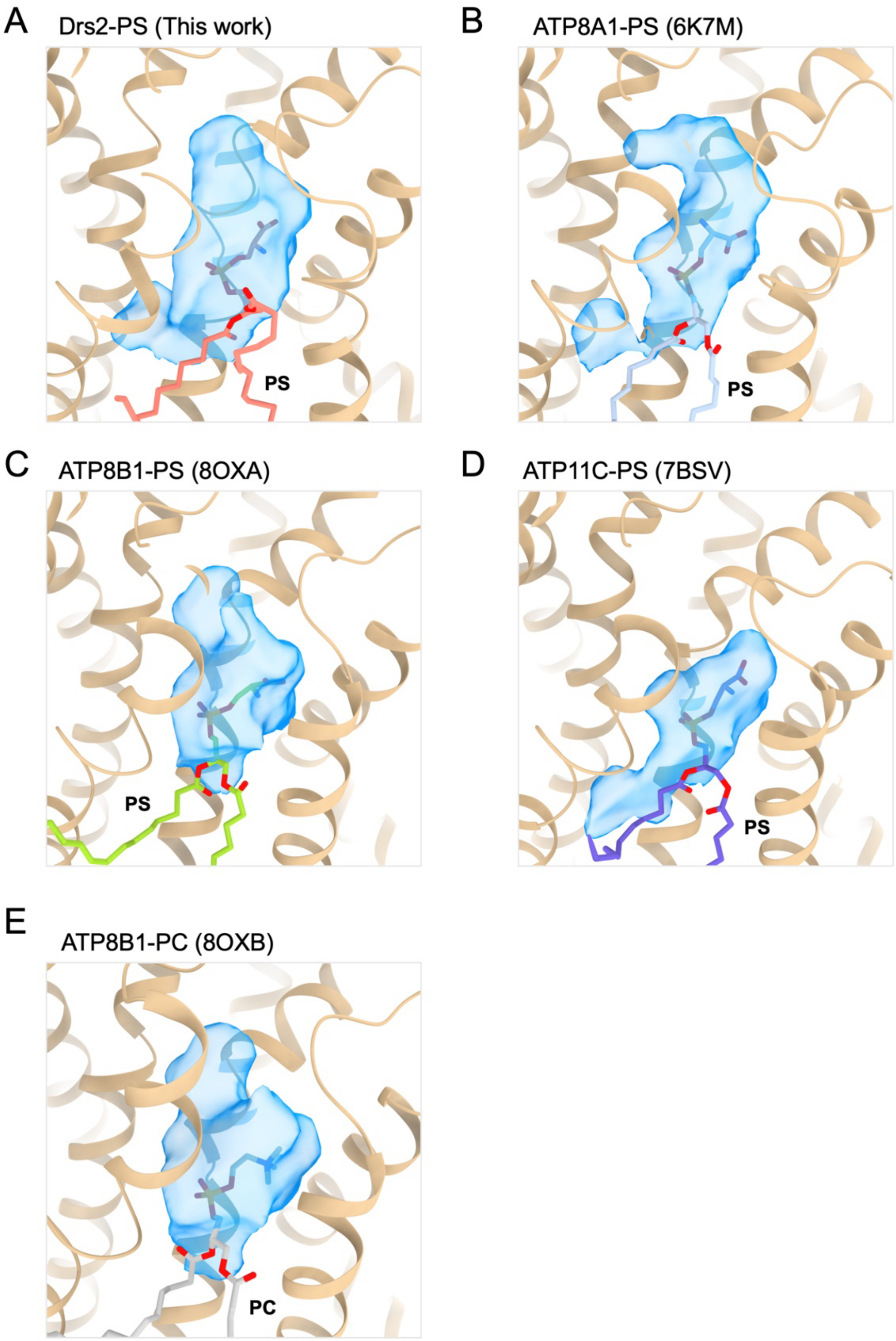

**Fig. S13: Comparison of lipid-occluded cavities**

Occluded lipid binding pocket of **(A)**, Drs2-PS, **(B)**, ATP8A1-PS, **(C)**, ATP8B1-PS, **(D)**, ATP11C-PS and **(E)**, ATP8B1-PC. Cavities were calculated by the pyKVFinder tool in ChimeraX-1.9 with a probe radius of 1.4 Å and displayed as a transparent blue surface (*78*). On comparison to Drs2, the binding pocket shape varies from being smaller and slimmer for ATP8A1 and ATP11C, to being similar for ATP8B1.

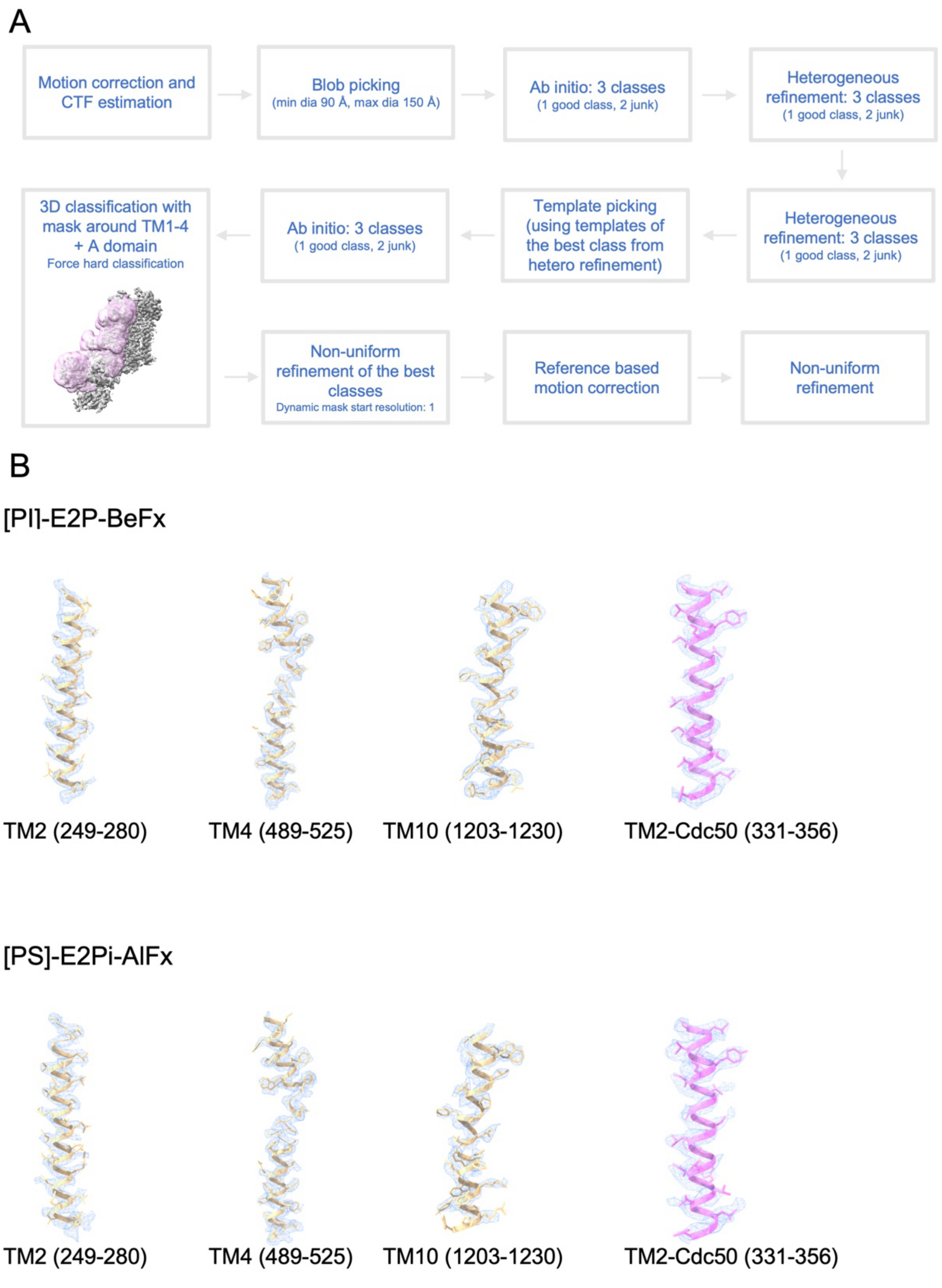

**Fig. S14: CryoSPARC data processing pipeline and representative cryo-EM maps for select TM segments**

**(A)**, Generic data processing pipeline performed in CryoSPARC for all datasets. The mask used in 3D classification is shown in pink. **(B)**, Cryo-EM maps and models of [PI]-E2P-BeFx and [PI]-E2Pi-AlFx showing the density of TM2, TM4, TM10 of Drs2 and TM2 of Cdc50.

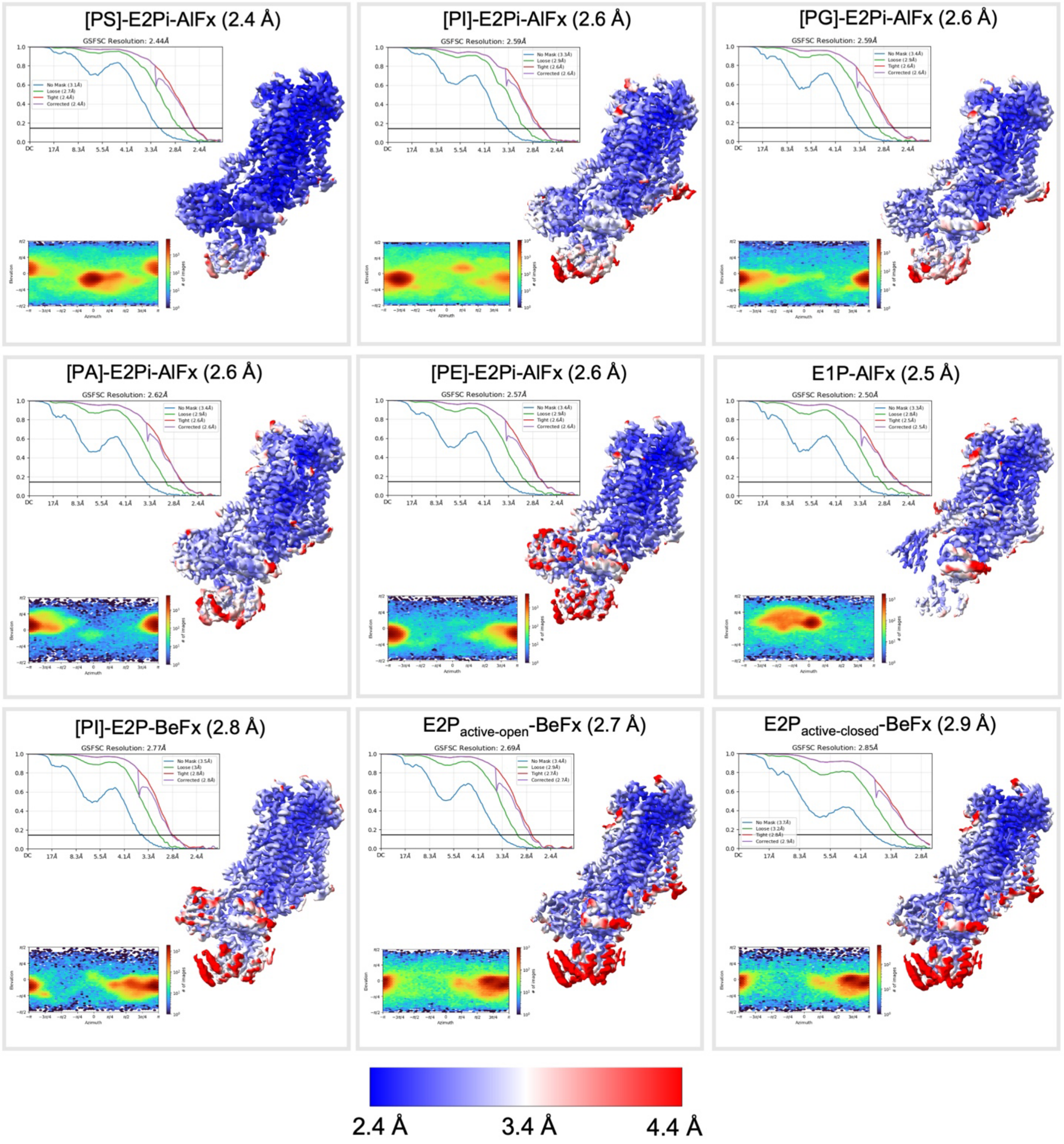

**Fig. S15: Local resolution, FSC and viewing distribution plots for all maps**

Each box shows the viewing distribution and Gold Standard Fourier Shell Resolution plot (FSC cut-off = 0.143), as well as the corresponding cryo-EM map colored according to local resolution. The resolution is given in a range from 2.4 Å (blue) to 4.4 Å (red).

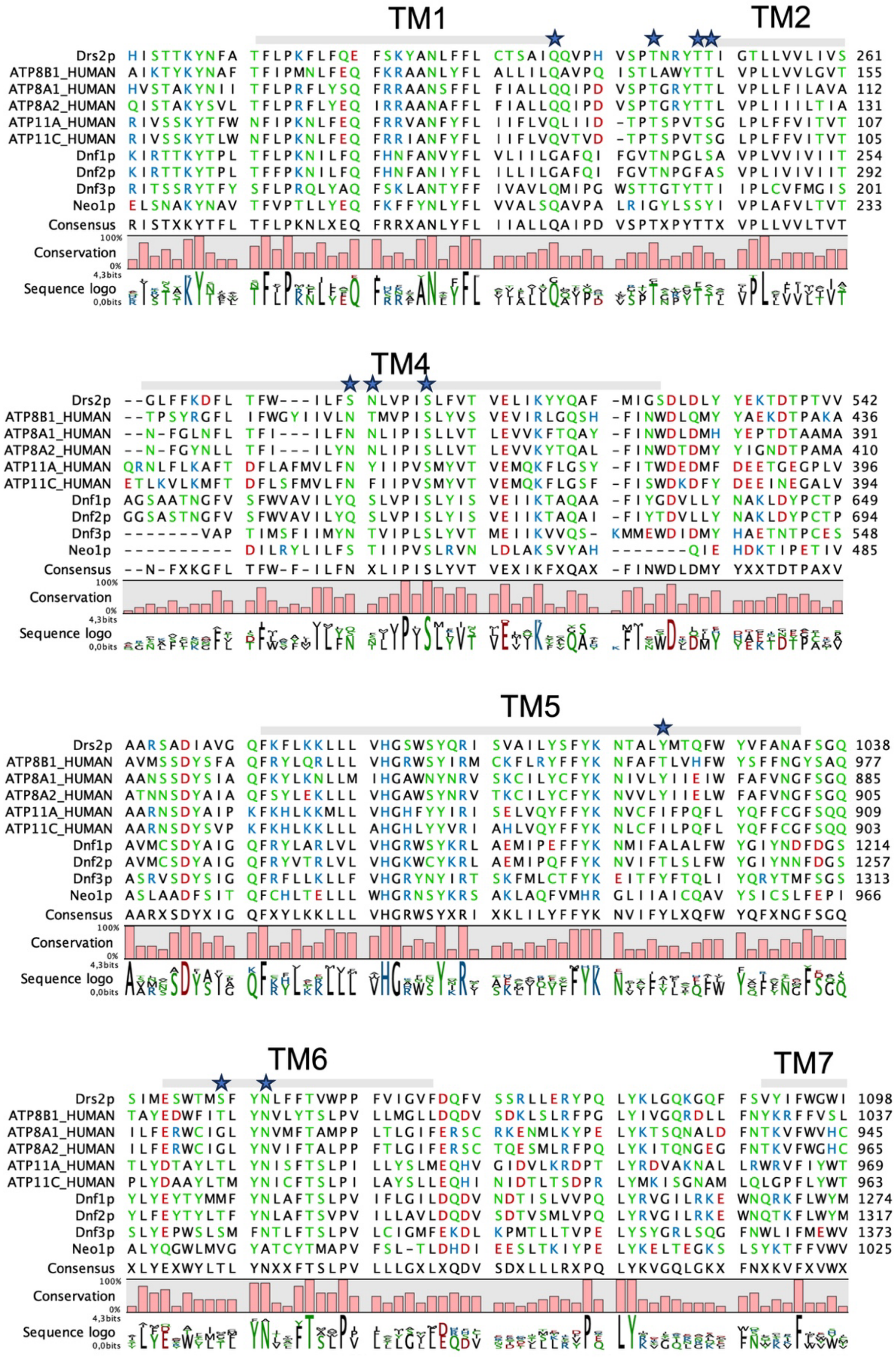
TM1
TM2
Drs2p HISTTKYNFA TFLPKFLFQE FSKYANLFFL CTSAIQQVPH VSPTNRYTTI GTLLVVLIVS 261
ATP8B1_HUMAN AIKTYKYNAF TFIPMNLFEQ FKRAANLYFL ALLILQAVPQ ISTLAWYTTL VPLLVVLGVT 155
ATP8A1_HUMAN HVSTAKYNII TFLPRFLYSQ FRRAANSFFL FIALLQQIPD VSPTGRYTTL VPLLFILAVA 112
ATP8A2_HUMAN QISTAKYSVL TFLPRFLYEQ IRRAANAFFL FIALLQQIPD VSPTGRYTTL VPLIIILTIA 131
ATP11A_HUMAN RIVSSKYTFW NFIPKNLFEQ FRRVANFYFL IIFLVQLIID -TPTSPVTSG LPLFFVITVT 107
ATP11C_HUMAN RIVSSKYTLW NFLPKNLFEQ FRRIANFYFL IIFLVQVTVD -TPTSPVTSG LPLFFVITVT 105
Dnf1p KIRTTKYTPL TFLPKNILFQ FHNFANVYFL VLIILGAFQI FGVTNPGLSA VPLVVIVIIT 254
Dnf2p KIRTTKYTPL TFFPKNILFQ FHNFANIYFL ILLILGAFQI FGVTNPGFAS VPLIVIVIIT 292
Dnf3p RITSSRYTFY SFLPRQLYAQ FSKLANTYFF IVAVLQMIPG WSTTGTYTTI IPLCVFMGIS 201
Neo1p ELSNAKYNAV TFVPTLLYEQ FKFFYNLYFL VVALSQAVPA LRIGYLSSYI VPLAFVLTVT 233
Consensus RISTXKYTFL TFLPKNLXEQ FRRXANLYFL IIALLQAIPD VSPTXPYTTX VPLLVVLTVT
100%
Conservation
0%
4,3bits
Sequence logo
0,0bits
TM4
Drs2p --GLFFKDFL TFW---ILFS NLVPISLFVT VELIKYYQAF -MIGSDLDLY YEKTDTPTVV 542
ATP8B1_HUMAN --TPSYRGFL IFWGYIIVLN TMVPISLYVS VEVIRLGQSH -FINWDLQMY YAEKDTPAKA 436
ATP8A1_HUMAN --N-FGLNFL TFI---ILFN NLIPISLLVT LEVVKFTQAY -FINWDLDMH YEPTDTAAMA 391
ATP8A2_HUMAN --N-FGYNLL TFI---ILYN NLIPISLLVT LEVVKYTQAL -FINWDTDMY YIGNDTPAMA 410
ATP11A_HUMAN QRNLFLKAFT DFLAFMVLFN YIIPVSMYVT VEMQKFLGSY -FITWDEDMF DEETGEGPLV 396
ATP11C_HUMAN ETLKVLKMFT DFLSFMVLFN FIIPVSMYVT VEMQKFLGSF -FISWDKDFY DEEINEGALV 394
Dnf1p AGSAATNGFV SFWVAVILYQ SLVPISLYIS VEIIKTAQAA -FIYGDVLLY NAKLDYPCTP 649
Dnf2p GGSASTNGFV SFWVAVILYQ SLVPISLYIS VEIIKTAQAI -FIYTDVLLY NAKLDYPCTP 694
Dnf3p -------VAP TIMSFIIMYN TVIPLSLYVT MEIIKVVQS- KMMEWDIDMY HAETNTPCES 548
Neo1p ---------- DILRYLILFS TIIPVSLRVN LDLAKSVYAH -------QIE HDKTIPETIV 485
Consensus --N-FXKGFL TFW-F-ILFN XLIPISLYVT VEXIKFXQAX -FINWDLDMY YXXTDTPAXV
100%
Conservation
0%
4,3bits
Sequence logo
0,0bits
TM5
Drs2p AARSADIAVG QFKFLKKLLL VHGSWSYQRI SVAILYSFYK NTALYMTQFW YVFANAFSGQ 1038
ATP8B1_HUMAN AVMSSDYSFA QFRYLQRLLL VHGRWSYIRM CKFLRYFFYK NFAFTLVHFW YSFFNGYSAQ 977
ATP8A1_HUMAN AANSSDYSIA QFKYLKNLLM IHGAWNYNRV SKCILYCFYK NIVLYIIEIW FAFVNGFSGQ 885
ATP8A2_HUMAN ATNNSDYAIA QFSYLEKLLL VHGAWSYNRV TKCILYCFYK NVVLYIIELW FAFVNGFSGQ 905
ATP11A_HUMAN AARNSDYAIP KFKHLKKMLL VHGHFYYIRI SELVQYFFYK NVCFIFPQFL YQFFCGFSQQ 909
ATP11C_HUMAN AARNSDYSVP KFKHLKKLLL AHGHLYYVRI AHLVQYFFYK NLCFILPQFL YQFFCGFSQQ 903
Dnf1p AVMCSDYAIG QFRYLARLVL VHGRWSYKRL AEMIPEFFYK NMIFALALFW YGIYNDFDGS 1214
Dnf2p AVMCSDYAIG QFRYVTRLVL VHGKWCYKRL AEMIPQFFYK NVIFTLSLFW YGIYNNFDGS 1257
Dnf3p ASRVSDYSIG QFRFLLKLLF VHGRYNYIRT SKFMLCTFYK EITFYFTQLI YQRYTMFSGS 1313
Neo1p ASLAADFSIT QFCHLTELLL WHGRNSYKRS AKLAQFVMHR GLIIAICQAV YSICSLFEPI 966
Consensus AARXSDYXIG QFXYLKKLLL VHGRWSYXRI XKLILYFFYK NVIFYLXQFW YQFXNGFSGQ
100%
Conservation
0%
4,3bits
Sequence logo
0,0bits
TM6
TM7
Drs2p SIMESWTMSF YNLFFTVWPP FVIGVFDQFV SSRLLERYPQ LYKLGQKGQF FSVYIFWGWI 1098
ATP8B1_HUMAN TAYEDWFITL YNVLYTSLPV LLMGLLDQDV SDKLSLRFPG LYIVGQRDLL FNYKRFFVSL 1037
ATP8A1_HUMAN ILFERWCIGL YNVMFTAMPP LTLGIFERSC RKENMLKYPE LYKTSQNALD FNTKVFWVHC 945
ATP8A2_HUMAN ILFERWCIGL YNVIFTALPP FTLGIFERSC TQESMLRFPQ LYKITQNGEG FNTKVFWGHC 965
ATP11A_HUMAN TLYDTAYLTL YNISFTSLPI LLYSLMEQHV GIDVLKRDPT LYRDVAKNAL LRWRVFIYWT 969
ATP11C_HUMAN PLYDAAYLTM YNICFTSLPI LAYSLLEQHI NIDTLTSDPR LYMKISGNAM LQLGPFLYWT 963
Dnf1p YLYEYTYMMF YNLAFTSLPV IFLGILDQDV NDTISLVVPQ LYRVGILRKE WNQRKFLWYM 1274
Dnf2p YLFEYTYLTF YNLAFTSVPV ILLAVLDQDV SDTVSMLVPQ LYRVGILRKE WNQTKFLWYM 1317
Dnf3p SLYEPWSLSM FNTLFTSLPV LCIGMFEKDL KPMTLLTVPE LYSYGRLSQG FNWLIFMEWV 1373
Neo1p ALYQGWLMVG YATCYTMAPV FSL-TLDHDI EESLTKIYPE LYKELTEGKS LSYKTFFVWV 1025
Consensus XLYEXWYLTL YNXXFTSLPV LLLGXLXQDV SDXLLLRXPQ LYKVGQLGKX FNXKVFXVWX
100%
Conservation
0%
4,3bits
Sequence logo
0,0bits

**Fig. S16: Sequence alignment**

Sequence alignment of selected flippases from human and yeast. The TMs are highlighted with a grey bar. The critical residues, which are involved in hydrogen bonding of the lipid substrate in the occluded state, are marked with a star. The residues are color-coded based on polarity.

| | [PS]-E2Pi-AlFx<br>EMDB-54729<br>PDB 9SBK | [PI]-E2Pi-AlFx<br>EMDB-54728<br>PDB 9SBJ | [PG]-E2Pi-AlFx<br>EMDB-54766<br>PDB 9SCQ |
|---|---|---|---|
| **Data collection and processing** | | | |
| Magnification | 130,000 x | | |
| Electron exposure (e-/Å) | 60 | | |
| Microscope | Titan Krios G3i | | |
| Camera | Gatan K3 | | |
| Defocus range (μm) | 0.8-1.8 | | |
| Pixel sixe (Å) | 0.647 | | |
| Symmetry imposed | C1 | | |
| Number of movies | 6,027 | 6,253 | 4,679 |
| Initial particle images (no.) | 1,942,142 | 2,738,689 | 909,414 |
| Final particle images (no.) | 674,835 | 830,151 | 318,826 |
| Map resolution<br>(FSC threshold) | 2.44<br>(0.143) | 2.59<br>(0.143) | 2.59<br>(0.143) |
| Map resolution range | 2.4-3.4 | 2.6-3.6 | 2.6-3.8 |
| **Refinement** | | | |
| Initial model used | 7OH6 | 7OH6 | 7OH6 |
| Model resolution (Å)<br>(FSC threshold) | 2.9<br>(0.5) | 3.1<br>(0.5) | 3.0<br>(0.5) |
| Map sharpening B factor($Å^2$) | 58.34 | 53.53 | 57.31 |
| Model composition<br>Non-hydrogen atoms<br>Protein residues<br>Ligands | <br>11,429<br>1,394<br>1 MG, 5 NAG, 1 ALF, 1 2Y5, 1 D39, 2 BMA, 1 MAN | <br>11,440<br>1,395<br>1 MG, 5 NAG, 1 ALF, 1 2Y5, 1 PIE, 2 BMA, 1 MAN | <br>11,433<br>1,395<br>1 MG, 5 NAG, 1 ALF, 1 2Y5, 1 DR9, 2 BMA, 1 MAN |
| B factors ($Å^2$, min/max/mean) | | | |
| Protein | 60.73/216.13/124.71 | 34.30/209.59/103.79 | 22.85/186.96/90.11 |
| Ligand | 81.80/165.13/122.35 | 50.72/133.79/94.07 | 45.65/109.11/79.03 |
| Water | 87.76/114.04/103.11 | 54.99/87.77/75.25 | 52.58/71.06/61.54 |
| R.m.s. deviations | | | |
| Bond lengths (Å) | 0.003 | 0.003 | 0.003 |
| Bond angles (°) | 0.622 | 0.686 | 0.646 |
| Validation | | | |
| MolProbity score | 1.52 | 1.51 | 1.47 |
| Clashscore | 5.86 | 7.16 | 7.38 |
| EMRinger | 1.61 | 1.54 | 1.25 |
| CaBLAM outliers | 1.09 | 0.73 | 0.44 |
| Ramachandran plot | | | |
| Favored (%) | 96.75 | 97.40 | 97.69 |
| Allowed (%) | 3.25 | 2.60 | 2.31 |
| Outliers (%) | 0.0 | 0.0 | 0.0 |

**Table S1: Data collection and refinement statistics**

| | [PE]-E2Pi-AlFx<br>EMDB-54725<br>PDB 9SBD | [PA]-E2Pi-AlFx<br>EMDB-54724<br>PDB 9SBC | E1P-AlFx<br>EMDB-54702<br>PDB 9SAV |
|---|---|---|---|
| **Data collection and processing** | | | |
| Magnification | 130,000 x | | |
| Electron exposure (e-/Å) | 60 | | |
| Microscope | Titan Krios G3i | | |
| Camera | Gatan K3 | | |
| Defocus range (μm) | 0.8-1.8 | | |
| Pixel sixe (Å) | 0.647 | | |
| Symmetry imposed | C1 | | |
| Number of movies | 4,296 | 5,950 | 1,040 |
| Initial particle images (no.) | 1,697,281 | 2,309,820 | 505,105 |
| Final particle images (no.) | 377,902 | 385,301 | 141,085 |
| Map resolution<br>(FSC threshold) | 2.57<br>(0.143) | 2.62<br>(0.143) | 2.51<br>(0.143) |
| Map resolution range | 2.6-3.9 | 2.6-3.7 | 2.5-3.5) |
| **Refinement** | | | |
| Initial model used | 7OH6 | 7OH6 | 7OH4 |
| Model resolution<br>(FSC threshold) | 2.9<br>(0.5) | 3.1<br>(0.5) | 2.8<br>(0.5) |
| Map sharpening B factor($Å^2$) | 72.75 | 75.51 | 65.41 |
| Model composition<br>Non-hydrogen atoms<br>Protein residues<br>Ligands | <br>11,437<br>1,395<br>1 MG, 5 NAG, 1 ALF, 1 2Y5, 1 PEE, 2 BMA, 1 MAN | <br>11,427<br>1,394<br>1 MG, 5 NAG, 1 ALF, 1 2Y5, 1 LIG, 2 BMA, 1 MAN | <br>8,193<br>992<br>1 MG, 5 NAG, 1 ALF, 1 2Y5, 1 BMA, 1 MAN |
| B factors ($Å^2$, min/max/mean) | | | |
| Protein | 13.19/155.15/62.38 | 22.33/169.70/82.97 | 42.59/179.52/93.89 |
| Ligand | 26.64/95.73/57.77 | 42.60/119.23/77.98 | 62.70/129.01/94.56 |
| Water | 28.90/58.36/40.57 | 45.73/79.73/63.26 | |
| R.m.s. deviations | | | |
| Bond lengths (Å) | 0.003 | 0.003 | 0.002 |
| Bond angles (°) | 0.674 | 0.628 | 0.655 |
| Validation | | | |
| MolProbity score | 1.61 | 1.63 | 1.20 |
| Clashscore | 8.95 | 7.59 | 3.59 |
| EMRinger | 2.17 | 2.09 | 3.40 |
| CaBLAM outliers (%) | 0.80 | 0.80 | 0.72 |
| Ramachandran plot | | | |
| Favored (%) | 97.33 | 96.68 | 97.76 |
| Allowed (%) | 2.67 | 3.32 | 2.24 |
| Outliers (%) | 0.0 | 0.0 | 0.0 |

**Table S2: Data collection and refinement statistics**

| | E2P$_{active-closed}$-BeFx<br>EMDB-54703<br>PDB 9SAW | E2P$_{active-open}$-BeFx<br>EMDB-54726<br>PDB 9SBH | [PI]-E2P-BeFx<br>EMDB-54727<br>PDB 9SBI |
|---|---|---|---|
| **Data collection and processing** | | | |
| Magnification | 130,000 x | | |
| Electron exposure (e-/Å) | 60 | | |
| Microscope | Titan Krios G3i | | |
| Camera | Gatan K3 | | |
| Defocus range (μm) | 0.8-1.8 | | |
| Pixel sixe (Å) | 0.647 | | |
| Symmetry imposed | C1 | | |
| Number of movies | 3,784 | | 5,730 |
| Initial particle images (no.) | 1,885,098 | | 3,419,145 |
| Final particle images (no.) | 79,125 | 150,303 | 162,484 |
| Map resolution<br>(FSC threshold) | 2.86<br>(0.143) | 2.69<br>(0.143) | 2.77<br>(0.143) |
| Map resolution range | 2.9-4.4 | 2.7-4.4 | 2.8-3.8 |
| **Refinement** | | | |
| Initial model used | 6ROJ | 6ROJ | 6ROJ |
| Model resolution<br>(FSC threshold) | 3.1<br>(0.5) | 2.9<br>(0.5) | 3.2<br>(0.5) |
| Map sharpening B factor($Å^2$) | -38.80 | 65.41 | 58.96 |
| Model composition<br>Non-hydrogen atoms<br>Protein residues<br>Ligands | <br>11,509<br>1,414<br>1 MG, 1 2Y5, 5 NAG, 1 BMA, 1 MAN | <br>11,525<br>1,414<br>1 MG, 1 2Y5, 5 NAG, 2 BMA, 1 MAN | <br>11,578<br>1,414<br>1 MG, 1 2Y5, 1 PIE, 5 NAG, 2 BMA, 1 MAN |
| B factors ($Å^2$, min/max/mean) | | | |
| Protein | 82.05/254.81/139.39 | 26.32/165.02/81.44 | 27.79/175.66/82.40 |
| Ligand | 92.12/186.07/141.28 | 44.32/110.20/78.73 | 40.12/126.89/74.86 |
| Water | | 55.70/79.42/66.97 | |
| R.m.s. deviations | | | |
| Bond lengths (Å) | 0.003 | 0.003 | 0.003 |
| Bond angles (°) | 0.638 | 0.606 | 0.687 |
| Validation | | | |
| MolProbity score | 1.43 | 1.41 | 1.59 |
| Clashscore | 7.26 | 5.04 | 9.19 |
| EMRinger | 2.14 | 2.46 | 1.56 |
| CaBLAM outliers | 0.85 | 0.92 | 1.35 |
| Ramachandran plot | | | |
| Favored (%) | 97.87 | 97.23 | 97.51 |
| Allowed (%) | 2.13 | 2.77 | 2.49 |
| Outliers (%) | 0.0 | 0.0 | 0.0 |

**Table S3: Data collection and refinement statistics**
